# The function of SIRT3 explored through the substrate interaction network

**DOI:** 10.1101/2020.12.10.414995

**Authors:** Jarmila Nahálková

## Abstract

SIRT3 is the mitochondrial protein lysine deacetylase with a prominent role in the maintenance of mitochondrial integrity vulnerable in the range of diseases. The present study examines the SIRT3 substrate interaction network for the identification of its biological functions in the cellular anti-aging mechanisms. The pathway enrichment, the protein function prediction, and the protein node prioritization analysis were performed based on 407 SIRT3 substrates, which were collected by the data mining. The substrates are interlinked by 1230 direct protein-protein interactions included in the GeneMania database. The analysis of the SIRT3 substrate interaction network highlighted Alzheimer’s disease (AD), Parkinson’s disease (PD), Huntington’s disease (HD), and non-alcoholic fatty liver disease (NAFLD) as the most associated with SIRT3 lysine deacetylase activity. The most important biological functions of SIRT3 substrates are within the respiratory electron transport chain, tricarboxylic acid cycle and fatty acid, triacylglycerol, and ketone body metabolism. In brown adipose tissue, SIRT3 activity contributes to the adaptive thermogenesis by the increase of energy production of the organisms. SIRT3 exhibits several modes of neuroprotective actions in the brain and liver including prevention of the mitochondrial damages due to the respiratory electron transfer chain failure, the quenching of ROS, the inhibition of the mitochondrial membrane potential loss, and the regulation of mitophagy. Related to its role in Alzheimer’s disease, SIRT3 activation performs as a repressor of BACE1 through SIRT3-LKB1-AMPK-CREB-PGC-1α-PPARG-BACE1 (SIRT3-BACE1) pathway, which was created based on the literature mining and by employing Wikipathways application. The pathway enrichment analysis of the extended interaction network of the SIRT3-BACE1 pathway nodes displayed the functional relation to the circadian clock, which also deteriorates during the progress of AD and it is the causative of AD, PD, and HD. The use of SIRT3 activators in combination with the stimulating effect of regular exercise is further discussed as an attractive option for the improvement of cognitive decline during aging and the progressive stages of neurodegeneration.

## Introduction

The reversion of the aging process or at least the postponing is a very ambitious task for the science of many generations. From the public health point of view, not the extension of the total length of the human age, but the improvement of the health and vitality of the aged people would be a desirable outcome of the research.

The sirtuin family is known for the significant functional link to aging and age-associated diseases. The experiments using SIRT3 knockout (KO) mice showed upregulation of most of the mitochondrial proteins, thus proving that SIRT3 is the major mitochondrial protein deacetylase [1]. The SIRT3 KO mouse model demonstrates the potential power of the SIRT3 in the protection against age-related diseases since mice develop a whole range of diseases including cancer, neurodegenerative, cardiovascular, and metabolic diseases [2]. SIRT3 crucially regulates mitochondrial bioenergetics, which is activated by nutrients and exercise [3]. The expression level of the SIRT3 declines in aging human stem cells (HSC-s) [4], the frontal lobe, and hippocampus of the old rats [5]. Intriguingly, the aging is reversible in the HSC-s just by the overexpression of SIRT3, which provides evidence of its biological functions in the cellular anti-aging mechanisms [4]. SIRT3 acts neuroprotective against mitochondrial disruption by the quenching of the reactive oxygen species (ROS), preventing the mitochondrial membrane potential loss, and as a sensor of the neurotoxic insults in the rat models of Alzheimer’s disease (AD) and the AD brain tissues [6]. The mentioned biological processes potentially mediating the exciting anti-aging function would be beneficial if they are utilized for the improvement of the health of the aging population.

The present study investigates the substrate interaction network of SIRT3, the NAD^+^-dependent protein lysine deacetylase with a major role in the protection of mitochondrial integrity, which is a susceptible target in several age-related diseases. The protein interaction network utilizes the hypothesis that the proteins interacting directly via protein-protein interactions are with high confidence participating in identical cellular and molecular functions [7]. Here it serves for the identification of the most important metabolic and signaling pathways navigating the biological effect of SIRT3 deacetylation activity. The use of pathway enrichment, protein function prediction, and protein node prioritization methods provides a new understanding of the anti-aging functions of the enzyme.

The substrates retrieved by the data mining yielded the interaction network, which involved SIRT3 and its 407 substrates linked by 1230 protein-protein interactions recognized by GeneMania application. The pathway enrichment analysis was performed by applying the GeneMania application run under the Cytoscape environment and complemented by the web-based STRING analysis. The protein node prioritization and the cluster analysis using Cyto-Hubba and MCODE applications were employed to determine the essential protein nodes and clusters with the major regulatory roles within the whole network. The results of the analysis are discussed in the frame of the mitochondrial functions of SIRT3 in age-related diseases such as metabolic syndrome, diabetes, cancer, and neurodegenerative diseases. The most important signaling and metabolic pathways linked to SIRT3 activity are highlighted for further regulatory options and therapeutical interventions.

## Materials and methods

### The list of SIRT3 substrates

The SIRT3 substrates were collected from the literature sources [8] [9] [10] [11] [12] [13] [14] [15] [16] [17] [18] [19] [20] [21] [22] [23] [24] [25] [26] [27] [28] [29] [30] [31] [32] (Tab. 1, Tab. 1S). One of the sources included a large-scale proteomic study, which identified the differentially acetylated liver mitochondrial proteins isolated from wild type (wt) and SIRT3 KO mice employing LC/MS-MS analysis. The SIRT3 substrates with the two-fold increase of the acetylation in the SIRT3 KO mice compared to wt mice and p<0.01 were selected for further analysis [30]. Moreover, there were used SIRT3 substrates from the proteomic analysis of the liver mitochondrial acetylome isolated from the wt and SIRT3 KO mice. The proteins with the minimum 2-fold acetylation increase in SIRT3 KO compared to wt mice and together with Welch’s t-test with Storey Correction <0,01 were selected for further analysis [31]. Finally, the SILAC studies of the differential protein acetylation from SIRT3 KO and wt murine embryonic fibroblasts and U2OS cells with SIRT3 upregulated by retroviral overexpression compared to the cells with shRNA silenced SIRT3 were also used as a source of the substrates. The differential threshold was set to the 2-fold acetylation change for the ratios SIRT3KO/wt and SIRT3KO/SIRT3 overexpression [32].

**Tab. 1.** The overview of the literature sources, which were used for the data mining of the SIRT3 substrates.

| Method/source | The references |
| --- | --- |
| Multiple analytical methods [8] | [9] [10] [11] [12] [13] [14] [15] [16] [17] [18] [19] [20]<br>[21] [22] [23] [24] [25] [26] [27] [28] [29] |
| LC/MS-MS; liver mitochondrial proteins; wild type (wt) *vs* SIRT3 KO mice | [30] |
| Nano reverse-phase LC/MS; liver mitochondrial proteins; wt mice *vs* SIRT3 KO | [31] |
| SILAC, LC/MS-MS; SIRT3 KO *vs* wt murine embryonic fibroblasts cells; U2OS cells with the upregulated SIRT3 *vs* cells with downregulated SIRT3 | [32] |

### GeneMania analysis

The protein names of SIRT3 substrates were converted to the GeneMania compatible symbols using the Protein Knowledgebase UniprotKB and the database OMIM^®^ (Online Mendelian Inheritance in Man^®^). GeneMania (3.5.2) [33–35] analysis was performed under Cytoscape (3.7.2) [36] environment utilizing the converted SIRT3 substrate list and including SIRT3 as a query (Tab. 1S). The analysis was run against the *H.sapiens* database (update 13-07-2017) and by including all types of interaction networks. The analysis was set up for the identification of the top 20 related genes and at the most 20 attributes by employing GO Molecular function weighting. The weighting method was selected based on the high probability of the sharing of the molecular functions between the SIRT3 substrates since the network of 408 proteins (including SIRT3) is interconnected by 1230 protein-protein interactions, which are included in the GeneMania database [7].

### String analysis

The web-based STRING analysis (v. 11) [37] was executed using the SIRT3 substrate list (Tab 1S) as a multiple protein query against the *H. sapiens* database. The settings were selected as follows: active interaction sources – text mining, experiments, and databases; minimum required interaction score – highest confidence (0.9); the maximum number of the interactors to show 1-st and 2-nd shell – none. The results of the functional enrichment analysis were exported with False Discovery Rate (FDR) value less or equal to 0,05. The interaction network was clustered by the Markov Cluster Algorithm (MCL) clustering method [38] with the inflation value equal to 10.

### The protein nodes prioritization

The high priority essential protein nodes and pathways of the SIRT3 substrate interaction network constructed by GeneMania were predicted by the Cyto-Hubba application (0.1) [39]. The application was set up for the use of the topological method Maximum click centrality (MCC), which provides according to the developers, the highest precision in the prediction of the essential nodes [39]. Further, the settings of the analysis were adjusted for the selection of the first-stage nodes and the display of the shortest path.

### Cluster analysis

Highly interconnected protein node clusters, which with high probability represent the functional complexes were identified by the Cytoscape application MCODE (v. 1.6) [40]. The following software settings were used for the analysis: Network scoring – Degree Cutoff: 2; Cluster finding – Haircut; Node score cutoff: 0.2; K-Core: 2; Max. Depth: 100.

### The pathway visualization

The pathway illustration was created employing the knowledgebase of the application WikiPathways [41] run under Cytoscape (3.7.2) [36] environment and by making use of the literature resources. The drawing was performed by Adobe Illustrator 2020 (24.2.1).

## Results

In the present study, an attractive age-related function of the main mitochondrial protein deacetylase SIRT3 is explored for the identification of the molecular mechanisms potentially applicable to therapeutical interventions. The attractive rejuvenation ability of SIRT3 is apparent in aged HSC-s, where it assists in cell survival under oxidative stress and renewing of the youthful phenotype [4]. Since SIRT3 KO mice display an astounding range of the diseases of aging [2] and due to the mentioned anti-aging effect of SIRT3 [4], the activators of SIRT3 and potentially also regulators of the functionally linked pathways are highly demanded for the therapeutical advancements. SIRT3 expression level in young cells is high, however, it decreases with the aging of the cells and tissues, which leads towards increased ROS and oxidative damages [42] [5]. These significant biological effects of SIRT3 are raising questions about the function of the interaction network linked to SIRT3 enzymatic activity, which could provide new regulatory options for therapeutical interventions.

The current study explores the protein interaction network of the substrates deacetylated by SIRT3, which employs the pathway enrichment, protein function prediction, protein node prioritization, and clustering methods. The analysis was applied for the identification of the most important diseases, along with signaling, and metabolic pathways linked to SIRT3 enzymatic activity. The protein node prioritization was employed to determine the essential protein nodes and clusters of the network with the major regulatory roles. Attention is also paid to the subcellular localization of the substrates processed by SIRT3, which assist in the additional interpretation of SIRT3 functionality in different cellular compartments.

### SIRT3 substrate interaction network

The current analysis utilizes the web-based STRING database as a tool for the construction and visualization of the interactive protein-protein interaction networks of SIRT3 substrates and the pathway enrichment analysis (Fig. 1, Tab. 2S). During the analysis is the protein dataset of SIRT3 substrates complemented with the known interactions from the public databases and through the computational predictions. The system further performs a classification based on the Gene Ontology, KEGG, high-throughput text mining, and hierarchical clustering of the network [43] [44][37], which provides useful outcomes for many biological questions.

**Fig. 1.**
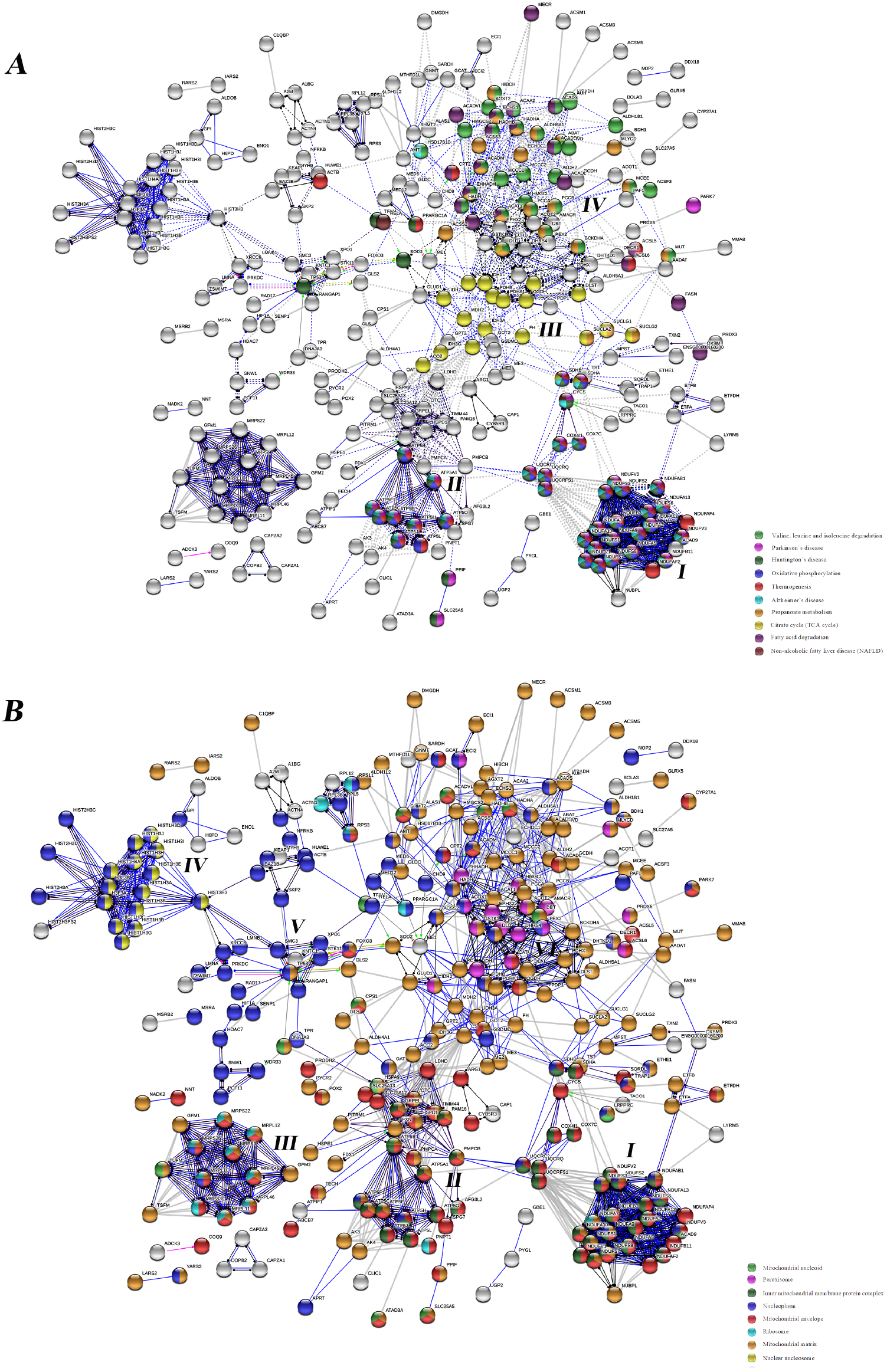
The pathway enrichment analysis of the SIRT3 substrate interaction network performed by the web-based STRING application. A. KEGG Pathways; B. GO Cellular Component. The analysis was performed using the SIRT3 substrate list as a multiple protein query against the *H. sapiens* database. The settings were selected as follows: active interaction sources – text mining, experiments, and databases; minimum required interaction score – highest confidence (0.9); the maximum number of the interactors to show 1-st and 2-nd shell – none.

The STRING database analysis is further complemented with the Cytoscape application GeneMania (Tab. 2), a consolidated pathway prediction analysis application based on the combined database annotations of genomic, proteomic, and mRNA expression data [45] [33] [34]. The software matches the query substrate list with the predicted and experimentally confirmed physical protein-protein interactions, genetic interactions, results of co-localization, and co-expression studies, and it fits the network with known signaling and metabolic pathways. One of the advantages of the GeneMania application is its integration with the Cytoscape environment, which allows the construction of the interaction network efficiently. The GeneMania database is continuously updated and expanded with additional data, which is not always the case with other types of interaction network platforms [35]. GeneMania analysis indicated that the most important pathways leading the functions of the SIRT3 substrate interaction network are the respiratory electron transport chain; general metabolic pathways; fatty acid, triacylglycerol, and ketone body metabolism, whereas AD and HD obtained lower scores (Tab. 2).

**Tab. 2.**
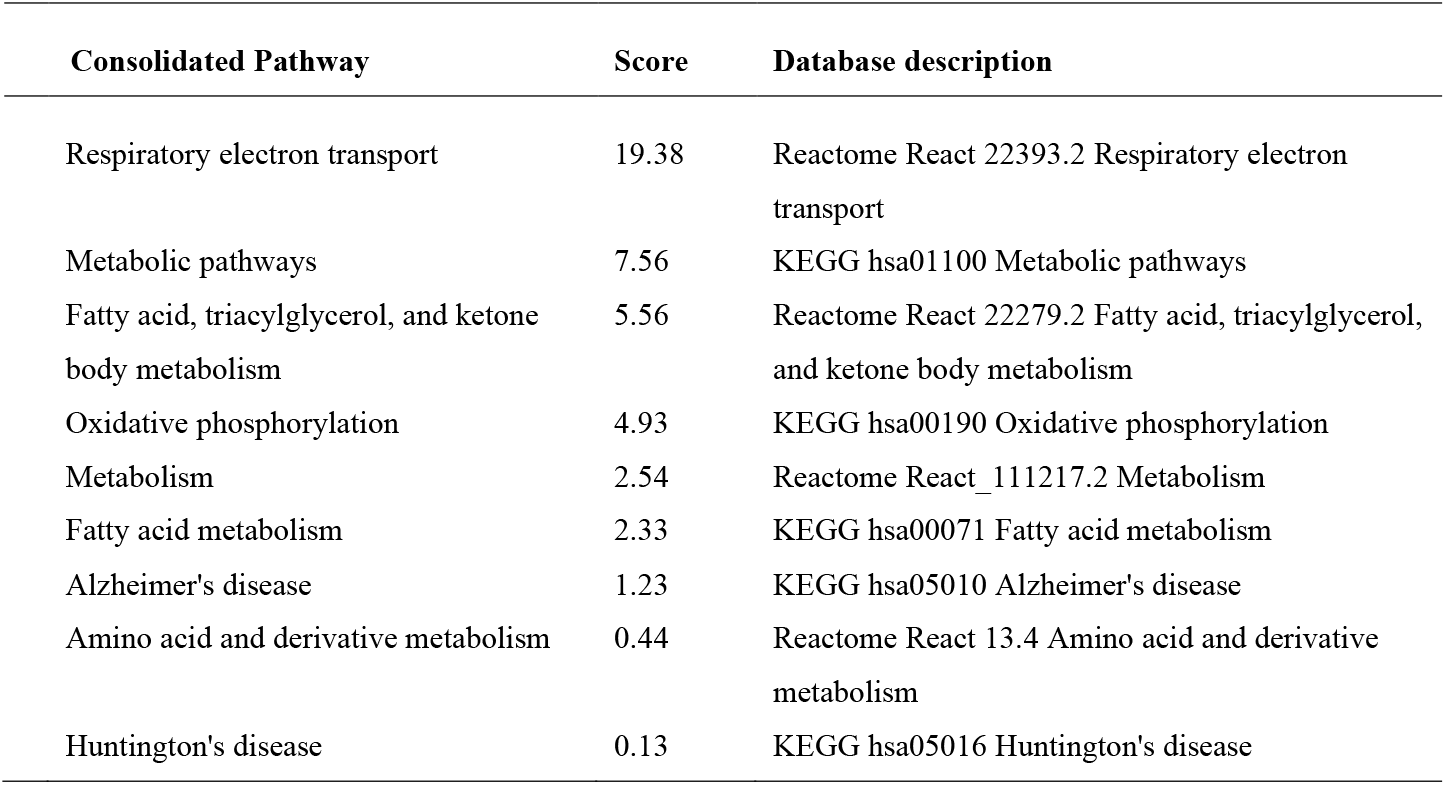
The result of the pathway enrichment analysis and the gene function prediction analysis of the SIRT3 substrate interaction network performed by the GeneMania application run under the Cytoscape (3.7.2) environment. The analysis was performed by utilizing the *H.sapiens* database (update 13-07-2017) and by including all types of interaction networks. The analysis was set up for the identification of the top 20 related genes and at the most 20 attributes using GO Molecular function weighting. *score (weight) expresses the predictive value, which GeneMania assigns to the pathways, how well they correspond to the query dataset compared to the non-query.

The additional results of STRING analysis using KEGG pathways classification visualized a linkage of SIRT3 substrates to Parkinson’s disease (PD); Huntington’s disease (HD); Alzheimer’s disease (AD); and non-alcoholic fatty liver disease (NAFLD), while creating discrete clusters by the MCL method (Fig. 1A) [38, 46]. SIRT3 expression is very high in the metabolically active tissues, including the brain, liver, heart, and brown adipose tissues, which correlates with functional roles in the neurodegenerative diseases pathologies, NAFLD, and the process of thermogenesis (Fig. 1A) [1].

The cluster I created by the MCL method contains the SIRT3 substrates associated with the oxidative phosphorylation. It consists of NADH: ubiquinone oxidoreductase (complex I) subunits, an enzyme that performs the electron transfer from NADH to the ubiquinone acceptor, a part of the respiratory electron transfer chain (Fig. 1A). The complex I activity is inhibited in SIRT3 KO mice, while the external SIRT3 can restore it, which suggests an activation effect through the deacetylase activity of SIRT3. The mRNA expression of the complex I subunits of the mitochondrial respiratory chain (MRC) also decreases in the brain of both early and definite AD patients compared to the control brains [47]. On the other hand, the expression of the complex III and IV subunits is enhanced, which might correspond to the increased energetic demands for ATP production in the AD brain [47]. ATP production is decreased >50 % in the hearts, kidneys, and livers of the SIRT3 KO mice, which further demonstrates the significant contribution of SIRT3 in cellular energy production [48].

The cluster also contains the complex II subunits of MRC, succinate dehydrogenase complex, subunit A (SDHA), and subunit B (SDHB), which are also associated with the mentioned 4 diseases (Fig.1A). The processing of the complex II subunits by SIRT3 was confirmed by some authors [49], while another report proved the deacetylation of SDHA by SIRT3 only with the partial stimulation effect on the activity of the whole complex II [50]. The limited regulation by acetylation makes sense since the complex II of MRC is an essential enzymatic complex and the complete inactivation is lethal [50]. The physical interaction of SIRT3 and SDHB was also demonstrated by two independent experiments, however without the detection of the specific acetylation sites [51]. The presence of the acetylation sites was demonstrated by the study used for data mining (Tab. 1, Tab. 1S) [30], where the acetylation of the lysines K160 and K169 revealed the highest increase due to SIRT3 KO [30]. However, the increase was lower than the two-fold threshold chosen for the present bioinformatic analysis.

Cluster II contains the protein substrates linked to AD, PD, and HD, which overlap with the functions in thermogenesis and oxidative phosphorylation (Fig. 1A). The cluster consists of the subunits of ATP synthase (complex V), the last enzyme of the mitochondrial oxidative phosphorylation chain, which produces the cell energy in the form of ATP by chemiosmotic proton efflux. The increased acetylation of the enzyme subunits occurs in the mitochondria of SIRT3 KO mouse livers and muscles, while the animals exhibit a lack of mitochondrial ATP production [3]. SIRT3 KO MEF cells also display significantly reduced ATP energy levels in mitochondria, which could be restored by the overexpression of wt SIRT3 but not by the inactive mutant [48]. SIRT3 stimulated by nutrients, calorie restriction (CR), and exercise deacetylates ATP-synthase subunits a, b, c, d, and OSCP, which leads to the increased mitochondrial energy supply and anti-aging effects [3]. Strikingly, SIRT3 is essential for the maintenance of the membrane potential of the healthy mitochondria, where it is bound to ATP synthase by pH and stress-sensitive bond [52]. The binding between ATP5O and SIRT3 can be disrupted by the change of the inner mitochondrial membrane potential, but not by the inhibition of the ATP production activity of ATP synthase [53]. The mitochondrial membrane potential is SIRT3 dependent, since after its disruption it recovers quickly in wt HeLa cells, but not in SIRT3 KO cells [53].

Cluster III consists of the tricarboxylic acid (TCA) cycle proteins, which according to the STRING are not associated with any of the major network-related diseases (Fig. 1A). However, the anaerobic metabolism of the glucose is very affected in the AD brain, where the most downregulated enzymes of the TCA cycle are located upstream of the succinyl-CoA [54]. SIRT3 substrates involved in cluster III (Fig. 1A; Tab. 2S) represent enzymes, which contribute to the dysregulated mitochondrial energetics during AD development. The most downregulated are the enzymes pyruvate dehydrogenase complex (PDHC; 41 %), isocitrate dehydrogenase complex (ICDH; 27 %), and α-ketoglutarate dehydrogenase complex (KGDHC; 57 %) [54]. Malate dehydrogenase (MDH) and succinate dehydrogenase (SDH) are significantly upregulated in the AD brain with 54 % and 44 % increase, respectively, while the rest of the enzymes involved in the TCA cycle remain unchanged. Interestingly, the clinical dementia rate (CDR) correlates well with the differential expression of the mentioned TCA cycle enzymes, while the best correlation was observed for PDHC [54].

Significantly, the decrease of PDHA1 and PDHC expression leads to the ATP deficits due to the metabolic switch towards glycolysis. PDHX, PDHB, PDHA1 (Fig. 1A, cluster III) are the subunits of PDHC, which is the first and irreversible step between the glycolysis and TCA cycle converting pyruvate to the acetyl-CoA. SIRT3 crucially activates the enzyme PDHA1 subunit by its deacetylation, which has an importance for the regulation of the Warburg effect in the cancer cells [55] [56]. Many cancer tissues, such as breast cancers contain inactivating mutation or gene deletion of SIRT3, which make favorable conditions for the growth of the glycolysis dependent cancer cells [57]. The limited level of PDHC causes the diverted production of acetyl CoA to oxaloacetate by the upregulated pyruvate carboxylase [58]. Due to the restricted metabolic pathway, the alternative energy is obtained by the reversal enzymatic reaction converting the oxaloacetate to succinate and by the electron transport chain of the complex I. The A*β* deposits of the AD brain cause the increased phosphorylation of PDHC leading to decreased energy outcome due to the metabolic switch from OXPHOS to glycolysis [58]. Age-related stimulation of pyruvate dehydrogenase kinase (PDK) deactivates PDHC activity through its phosphorylation and the resulting ATP production deficits [59] lead towards mitochondrial damage and synaptic deterioration.

Cluster IV is formed by SIRT3 substrates involved in fatty acid degradation and the branched-chain amino acid metabolism (Fig. 1A). The increased acetylation of PDH at SIRT3 KO mice leads to a metabolic switch by increased fatty acid oxidation and accumulation of the lactate occurring in muscles [24]. Further, it causes hyperacetylation of the long-chain Acyl-CoA dehydrogenase, which causes an accumulation of the long-chain fatty acid metabolic intermediates. The defect in fatty acid oxidation is further exacerbated by CR in SIRT3 KO compared to wt mice [19].

Conclusively, the result of the analysis emphasizes the crucial role of SIRT3 in the mitochondrial ATP production using chemiosmotic proton gradient, metabolism of the glucose degradation, and fatty acid oxidation. The processing of the SIRT3 substrates is primarily related to the brain related neurodegenerative diseases AD, PD, HD, liver associated NAFLD, and the process of the thermogenesis occurring in the brown adipose tissue.

### The protein node prioritization analysis

The SIRT3 substrate interaction network constructed by GeneMania was further subjected to the protein node prioritization using the Cyto-Hubba application (Fig. 2). The analysis revealed the most essential protein nodes possibly representing the pharmaceutical targets of the interaction network.

**Fig. 2.**
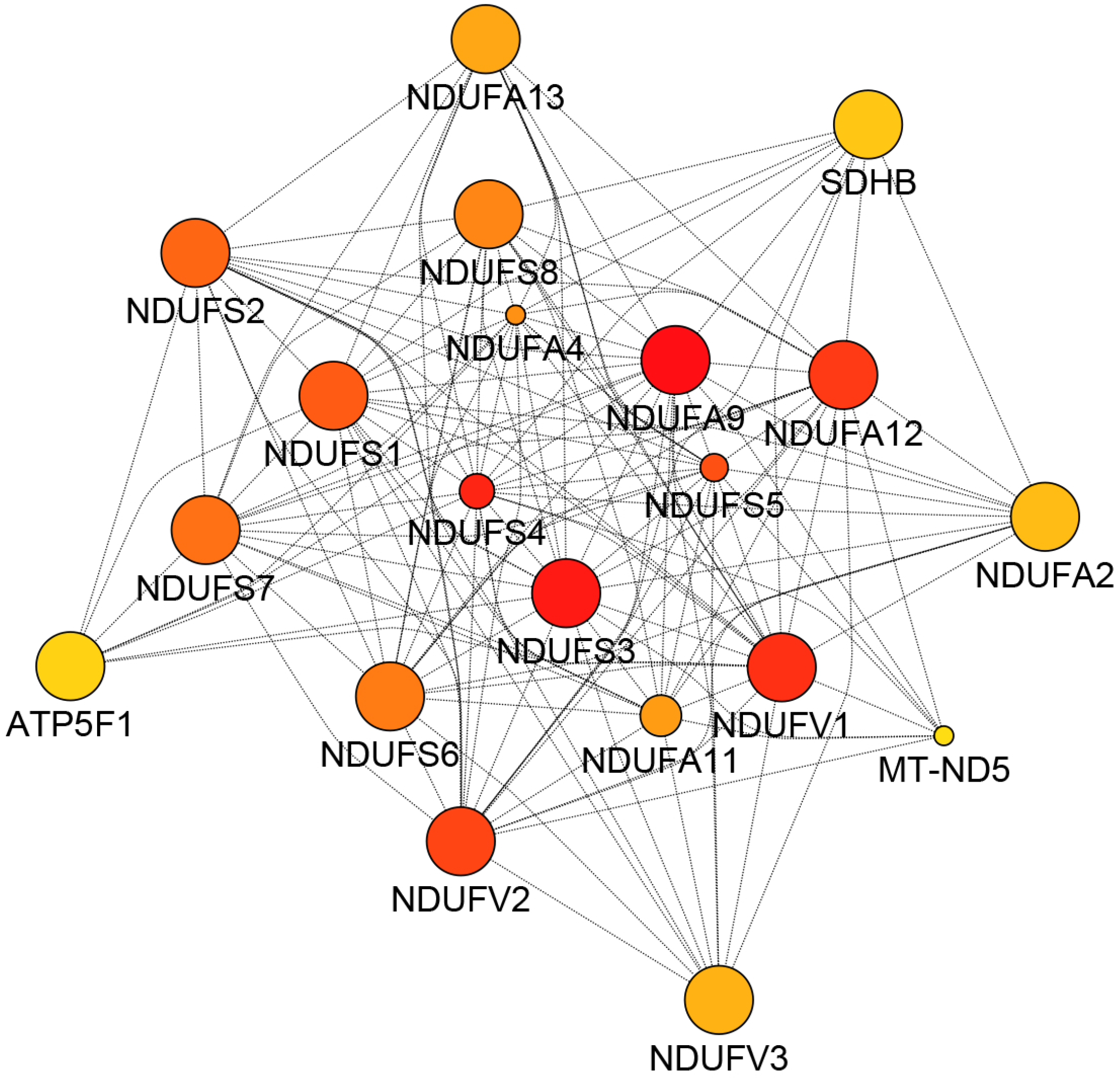
The protein node prioritization of the SIRT3 substrate interaction network predicted by the cytoHubba (0.1) application. The network was constructed by applying the GeneMania (3.5.2) application run under Cytoscape (3.7.2) environment utilizing the *H.sapiens* database (update 13-07-2017) and including physical interactions only. The analysis was set up for the identification of the top 20 related genes and at the most 20 attributes using GO Molecular function weighting. The cytoHubba analysis was adjusted for the use of the topological method Maximum click centrality (MCC), and the display of the shortest path. The proteins are shown from the highest (red) to the lowest (yellow) ranked nodes.

The Cytoscape application Cyto-Hubba identified the subunits of NADH: ubiquinone oxidoreductase, representing the complex I of MRC among the nodes with the highest rank (Fig. 2), which suggests that they have the most important regulatory effect within the entire SIRT3 substrate interaction network (Tab. 2). Additional high-ranked nodes are SDHB, succinate dehydrogenase subunit B, complex II; and ATPF1, which encodes the subunit B of ATP synthase Fo unit, which belongs to the complex V of MRC.

The MCODE cluster analysis further identified the most important interaction subgroups. The MCODE analysis prioritized the subnetwork of 70 protein nodes linked by 3913 interactions (score 63, 043) from the network of SIRT3 substrates extended by GeneMania application. The priority subnetwork included the protein nodes involved in the oxidative phosphorylation, respiratory transport chain, and the interaction networks of the diseases AD and HD (data not shown), which defines the main targets of the SIRT3 deacetylase activity.

As a result, the protein node prioritization determined 20 the most essential proteins of the SIRT3 substrate interaction network, which are mainly the subunits of the complex I, complex II, and complex V of MRC. The complementary prioritization method also confirmed that the mitochondrial bioenergy production through oxidative phosphorylation and MRC has the highest rank within the interaction network of 407 substrates.

### The subcellular localization of SIRT3 and its substrates

Mostly described as a mitochondrial protein deacetylase, there exists disagreement about the subcellular location of SIRT3 in other compartments debated by different authors. According to some reports, SIRT3 shuttles between the nucleus and mitochondria, which occurs upon exposure to a cellular stressor or after overexpression. SIRT3 is exported to the mitochondria after processing by the cleavage of N-terminal 142 aa residues [60].

The nuclear localization of SIRT3 was, though highly doubted by some authors since the deacetylase is according to the independent experiments located exclusively in the mitochondria [61] [62]. Contrary to this observation, the localization of SIRT3 in the nucleus was suggested by the immunofluorescence, when the nuclear signal was detected on Z-stack by confocal microscopy, despite much weaker than in the mitochondria. The deacetylation of exclusively nuclear substrates also confirms the existence of nuclear SIRT3. Among the sirtuins, SIRT3 has the best ability to remove *β*-hydroxybutyryl (bhb) from the nuclear substrate H3K9bhb [63]. An additional nuclear substrate of SIRT3 is H3K56 [64]. However, if the deacetylation of mitochondrial substrates encoded in the nucleus occurs before or after the export from the nucleus remains to be determined.

In the present study, SIRT3 substrates used for data mining originated from the analysis of the mitochondrial fractions [30][31] with the major overrepresentation of the mitochondrial proteins. The exception is the SILAC study [32] (Tab. 1), which used the whole-cell lysis step before LC-MS/MS identification. The SIRT3 substrate list obtained from this study indeed contains the multiple histone proteins (Fig. 1B), while their occurrence on lists of the substrates from other sources is limited only to Hist1h4a [30][31]. For the visualization of the subcellular localization of the collected SIRT3 substrates, the interaction network was processed through the GO Cellular Component classification. The analysis exhibited four main subcellular compartments nucleus, mitochondria, cytoplasm, and peroxisomes as the location of SIRT3 substrates (Fig. 1B). The majority of SIRT3 substrates co-localized also in the cytoplasm (data not shown) excluding the histone proteins.

Cluster I and cluster II contains the SIRT3 substrates located both in the mitochondrial compartments and the nucleus (Fig. 1B). The complex I respiratory electron transport chain subunits (NDUFS1-3, NDUFS6), acyl-CoA dehydrogenase subunit 9 (ACAD9), and NDUFAB1 are located both in the mitochondria and the nucleoplasm. ATP synthase peripheral stalk membrane subunit B (ATP5F1) is also localized in the nucleoplasm, the mitochondrial membrane, and the mitochondrial matrix (Fig. 1B). PNPT1 involved in the processing of the mitochondrial mRNA and polyadenylation localizes in mitochondrial ribosomes and the mitochondrial envelope [65]. SLC25A13, a mitochondrial calcium-binding carrier, transports the cytoplasmic glutamate for the mitochondrial aspartate through the inner mitochondrial membrane [66], locates in the mitochondrial matrix, and mitochondrial nucleoid (Fig. 1B).

Cluster III contains mitochondrial proteins encoded in the nucleus and localized in the different mitochondrial compartments (Fig. 1B). SIRT3 substrates GFM1, GFM2, and TSFM included in these clusters are the mitochondrial elongation factors important for the protein translation in the mitochondria and they are localized in the mitochondrial matrix. Their mutations are causative of the mitochondrial diseases due to MRC deficiencies [67] [68] [69]. Another translation elongation factor TUFM [70] is localized both in the matrix and nucleoid of the mitochondria (Fig. 1B). SIRT3 was demonstrated to regulate the translational activity of the mitochondrial ribosomes [18]. The mitochondrial ribosomal proteins MRPL10, MRPL11, MRPL12, MRPL45, MRPL46, MRPS22, MRPS26, MRPS30, and MRPS9 involved in this cluster are encoded by the nuclear DNA.

Cluster IV contains the nuclear histone proteins and several exclusively nucleoplasmic proteins. The enzymatically active SIRT3 fulfills the function in the chromatin silencing by the deacetylation of H4K16Ac and H3K9Ac [60] and H3K56Ac [62].

Cluster V contains SIRT3 substrates located in the nucleosomes and cluster VI is defined by the unique peroxisome localization and the cellular functions in *β*-oxidation of the fatty acids and the inactivation of ROS. It is possible, that the substrates are firstly processed in mitochondria and nuclei before they relocate to the peroxisomes. This observation is validated by the report of existing membrane contact sites of peroxisome membranes with other organelles including mitochondria and the nuclei [71].

As a conclusion, the collected SIRT3 substrates were sorted into distinct clusters based on their subcellular localization in the mitochondrial, nuclear, and peroxisomal subcellular compartments. Besides the cluster of the solely nuclear histone substrates, other nuclear-encoded proteins with significant functions in the mitochondria are deacetylated by SIRT3 probably after their export from the nucleus. This claim, however, will need the final experimental clarification together with the experimental proof of the occurrence of SIRT3 in peroxisomes. Since SIRT3 was not previously detected in the peroxisomes, but the peroxisomal membrane fusion with other organelles might be a feasible explanation of the transport mechanism of the processed substrates into the peroxisomes.

### The metabolic syndrome and diabetes

The GeneMania analysis visualized multiple cellular metabolic functions of the SIRT3 substrate interaction network, which highlighted its roles in the primary metabolism of carbohydrates, fatty acids, and amino acids (Tab. 2). The metabolic functions of SIRT3 can be illustrated on the observations made in SIRT3 KO mice fed a high-fat diet. The animals exhibit the downregulation of the liver SIRT3, which further increases acetylation of the mitochondrial proteins, accelerate obesity, the insulin-resistance, and metabolic syndrome [72]. SIRT3 is also downregulated in the muscles of the diabetes mouse model, which is related to its metabolism-related functions [73].

SIRT3 expression and its mitochondrial function are closely related to metabolic syndrome, and dementia. SIRT3 deficiency can cause the formation of the inflammasomes in the brain of the metabolic syndrome patients due to mitochondrial dysfunction, which is further causative of the cognitive decline [74]. The middle-age metabolic syndrome, type 2 diabetes, and overweight are also a precondition for the onset of dementia, AD, and vascular dementia at a late age [75] [76]. The activation of SIRT3 is a promising approach for the prevention of metabolic diseases accompanied by the risk of the development of dementia.

### The regulation of BACE1

Related to AD pathology, SIRT3 enzymatic activity interestingly inhibits the A*β* production in the brain through the regulation of *β*-secretase (BACE1), an enzyme of the first and the rate-limiting step in the APP processing. The pathway mediating the regulatory effect of SIRT3 on the activity of BACE1 was constructed utilizing the literature mining, WikiPathways, and GeneMania application (Fig. 3, Fig. 4, Tab. 3).

**Fig. 3.**
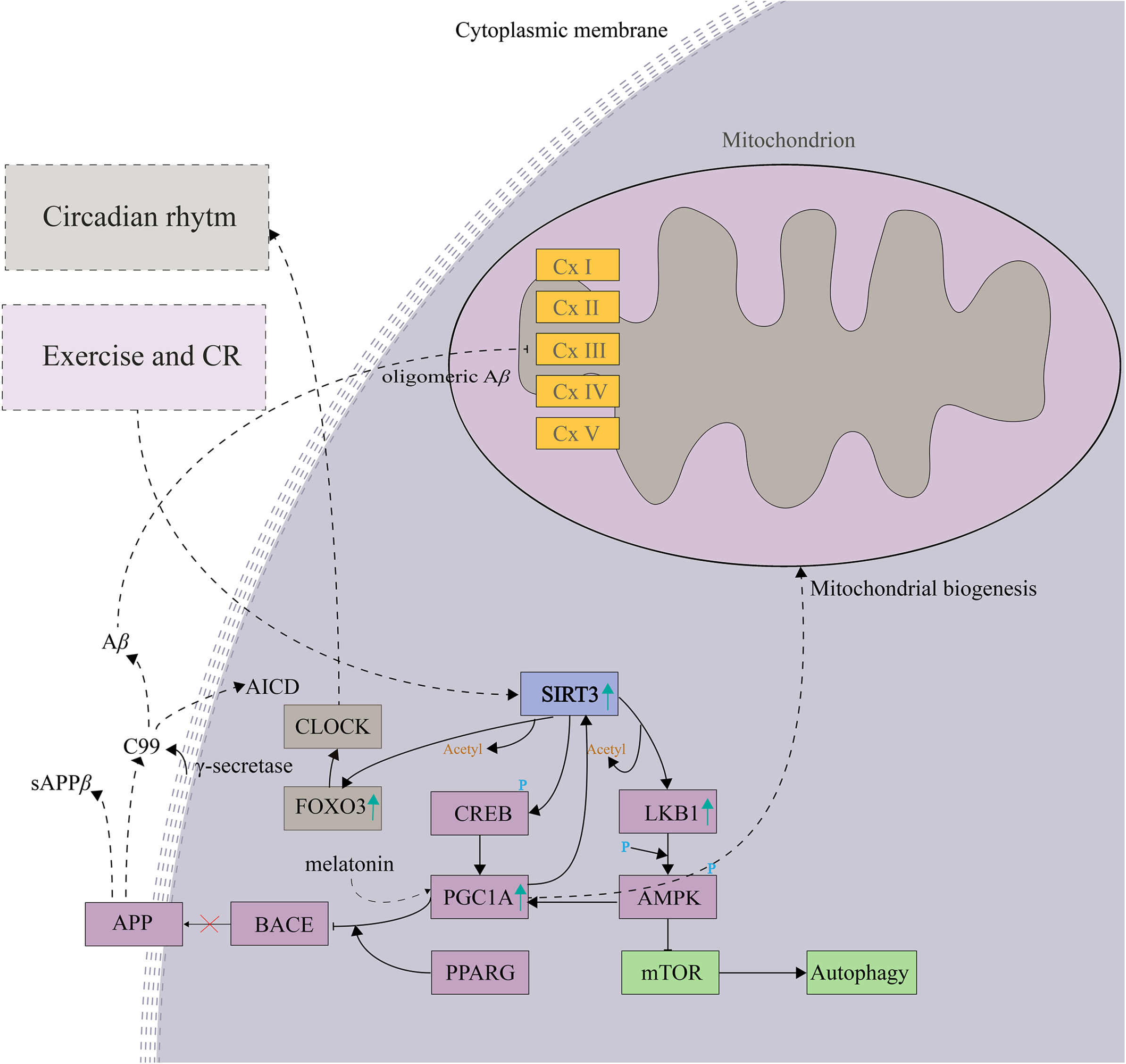
The enzymatic activity of SIRT3 inhibits the A*β* production in the brain by the regulation of *β*-secretase (BACE1) through the SIRT3-LKB1-AMPK-CREB-PGC-1α-PPARG-BACE1 pathway.

**Fig. 4.**
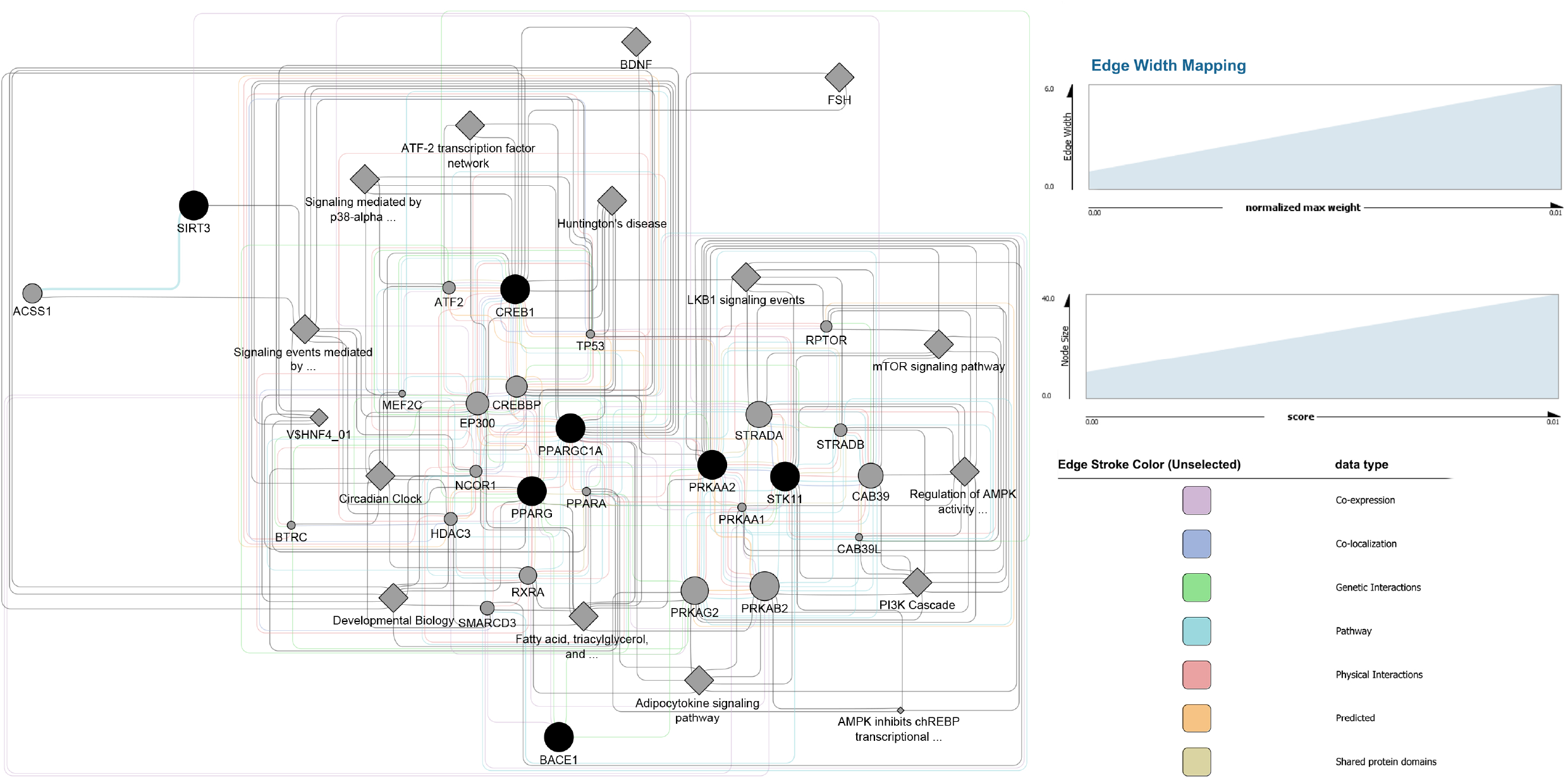
The protein interaction network constructed by GeneMania (3.7.2) where all proteins of the pathway SIRT3-LKB1-AMPK-CREB-PGC-1α-PPARG-BACE1 were used as a query. The analysis was run against the *H.sapiens* database (update 13-07-2017) and by including all types of interaction networks. The analysis was set up for the identification of the top 20 related genes and at the most 20 attributes using GO Molecular function weighting. The result showed the Circadian Clock as a pathway with the highest score (weight) indicating the common functions with the regulation of BACE1. * HDAC class I and III

**Tab. 3.**
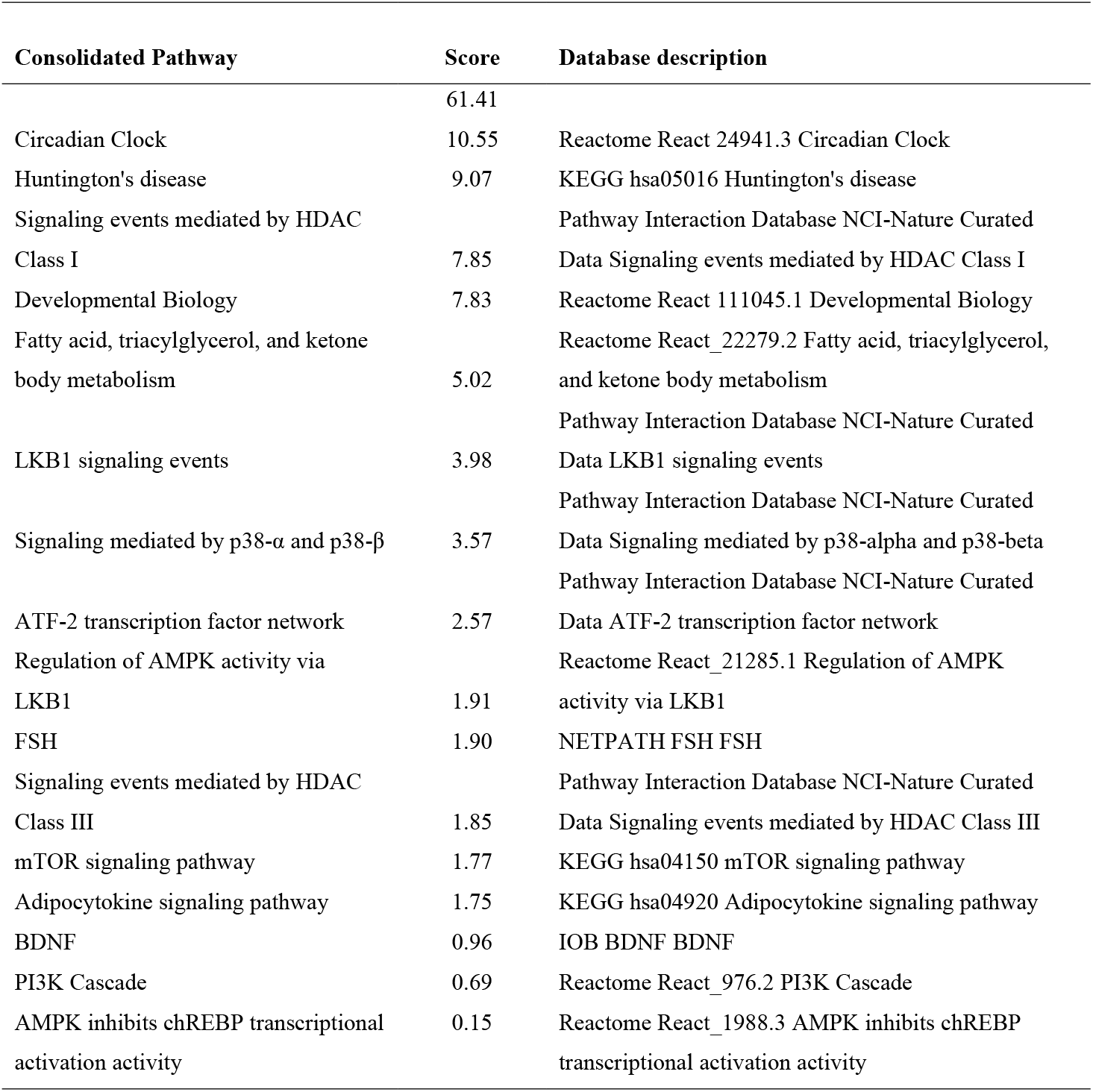
The result of the pathway enrichment analysis performed by GeneMania application run under the Cytoscape (3.7.2) environment, where all proteins of the pathway SIRT3-LKB1-AMPK-CREB-PGC-1α-PPARG-BACE1 were used as an input. The analysis was run against the *H.sapiens* database (update 13-07-2017) and by including all types of interaction networks. The analysis was set up for the identification of the top 20 related genes and at the most 20 attributes using GO Molecular function weighting. The circadian clock pathway obtained the highest score in the analysis. *score (weight) expresses the predictive value, which GeneMania assigns to the pathways, how well they correspond to the query dataset compared to the non-query.

SIRT3 activated by the exercise and/or CR deacetylates and activates serine/threonine-protein kinase 11 (LKB1, STK11; Table 1S), which is followed by the phosphorylation of AMPK and the activation of the pathway SIRT3-LKB1-AMPK [77]. It results in the upregulation of PGC-1*α* [78], a repressor of BACE1, which finally inhibits the production of A*β* in the brain [79] (Fig. 3). Alternatively, SIRT3 also activates CREB through its phosphorylation, which stimulates the PGC-1α promoter directly [81]. Further PGC-1α requires the presence of PPARG for its effect on the expression of BACE1. In agreement with this model, the expression level of PGC-1α is decreased in the hippocampus of AD patients compared to the age-matched controls [80], which supports the pathological significance of the SIRT3 - LKB1 - AMPK - CREB - PGC-1*α* – PPARG – BACE1 pathway (Fig. 3).

SIRT3 is a negative regulator of autophagy, which is involved in the proteolytic degradation of the protein aggregates and the whole organelles and it is often impaired in AD [82]. The specific overexpression of SIRT3 in the mouse liver and hepatocytes causes AMPK and autophagy suppression and oppositely, the downregulation of SIRT3 activates autophagy in both the liver-specific knockdown and hepatocytes [83].

The age-related decrease of PGC-1α correlates with increased expression of BACE1 and with the pathological observations of the neuritic *β*-amyloid plaques in AD brains. The viral overexpression of PGC-1α in the neuronal culture significantly decreases the production of the extracellular A*β*42, which confirms that the regulation of PGC-1α is an effective target for AD therapy [84]. PGC-1*α* plays a role in mitochondrial bioenergetics by the stimulatory effect on the mitochondrial biogenesis, the oxidative phosphorylation, and by the cell-specific upregulation of the uncoupling proteins UCP-s. The promotion of the uncoupled mitochondrial respiration has an importance in the adaptation to the cold temperature and the prevention of obesity [85]. In the nucleus PGC-1*α* directly promotes the transcription of the nuclear-encoded MRC subunits such as cytochrome c, *β*-ATP synthetase, COXIV, and co-activates the main mitochondrial transcription factor mtTFA [85].

With the purpose to identify the biological functions, which correlate with the SIRT3-LKB1-AMPK-CREB-PGC-1α-PPARG-BACE1 pathway (Fig. 3) [77][78][79][80][81], the pathway enrichment analysis was performed employing GeneMania application (Fig. 4) by applying the application compatible protein names SIRT3, LKB1, AMPK, CREB1, PGC1A, PPARG, and BACE1 as an input. Interestingly, the highest weight in the pathway enrichment exhibited the Circadian Clock (Fig 4, Table 3). As a common feature, PGC-1*α* also plays a central role in the regulation of the key oscillators of the circadian rhythm BMAL1 and CLOCK, which coordinate functions of the mammalian clock and energy metabolism [86]. The circadian rhythm regulator melatonin activates PGC-1α, which further interacts with estrogen-related receptor α (ERRα) directly bound on the SIRT3 promoter and enhances its transcription activity [87].

The constructed pathway demonstrating the mode of the action of SIRT3 on BACE1 activity is potentially interesting for AD treatment development. The pathway provides an overview of the mutual regulatory effect of the protein nodes which could be potentially attractive for multitarget therapy development.

### AD and the Circadian Clock

The Circadian Clock pathway significantly enriched by the interaction network analysis of the SIRT3-LKB1-AMPK-CREB-PGC-1α-PPARG-BACE1 pathway (Fig. 4, Tab. 3) is also substantially altered in AD. The rhythmic expression of the circadian oscillators BMAL1, PER1, and PER2 differs in the AD pineal gland, cingulate cortex, and bed nucleus of stria terminalis compared to the control [88]. The correlation study of the circadian clock function with the metabolic disturbances, the involvement of ROS, and autophagy leads to the conclusion that the disruption of the circadian clock can be the causative of AD, PD, and HD in humans [89]. The circadian clock also regulates the rhythm of NAD^+^-dependent SIRT3 through the nicotinamide riboside (NR) pathway, which is activated by the nutrient availability during the feeding period [90].

Interestingly, the acetylation of cellular proteins also exhibits diurnal patterns since most of the nuclear and cytoplasmic proteins are acetylated during the night and mitochondrial proteins are modified during the day. The rhythmic acetylation is functionally linked to the rhythmic deacetylation, while SIRT3 is the main mitochondrial protein deacetylase [90].

The regulation of the circadian cycle occurs through the SIRT3-FOXO3-CLOCK pathway (Fig. 3), where FOXO3 is the crucial regulator of the circadian cycle. At the conditions of low insulin occurring in the aging brain [91], it binds directly to the CLOCK promoter and regulates its transcription [92]. The activation of SIRT3 increases FOXO3 expression through their direct physical interaction, which, however, does not depend on SIRT3 deacetylase activity [93] [94]. In a conclusion, the function of the pathway SIRT3-LKB1-AMPK-CREB-PGC-1α-ERRA-PPARG-BACE1 is closely related to the circadian clock pathway, which deteriorates in AD and it appears to be the causative of the disease. SIRT3-FOXO3-CLOCK pathway is linked to metabolic disruption, ROS production, and autophagy, which also have an importance in AD pathology.

### The regulation of tau pathology

The deacetylation of tau by SIRT3 plays a significant neuroprotective role in AD. The cognitive impairment in AD correlates with the tau acetylation at K274 and K283, which substantially upregulates by a high A*β* level when compared to the control brain [97]. AD brain tissues also contain specific tau acetylation at K280, while the modification was identified as a reason for the pathological tau aggregation [98]. Stunningly, the neuroprotection is established just by the increased SIRT3 expression, which reduces the level of the total, and the acetylated tau in the A*β*42 pretreated primary neurons from the transgenic mice expressing human tau [99] and in the hippocampal cells transfected by Sirt3 cDNA compared to SIRT3 shRNA [100]. Therefore, the possibility to regulate the tau pathology by the SIRT3 activators seems a hopeful strategy, especially if the natural products would be used.

### TP53 and AD

An additional aspect of SIRT3 multifunctionality is the specific regulation of the important substrate p53 through its deacetylation (Fig. 1; Table 1S) implicated in AD pathology.

In the neurons of AD patients, the expression level of p53 and its K320 acetylated form increases both in the nuclear and mitochondrial fractions. At the same time SIRT3 expression decreases, which is associated with neuronal damage and mitochondrial dysfunction [101].

The upregulation of p53 in the brain induces cell necrosis in the presence of oxidative stress. P53 physically interacts with the critical regulator of the mitochondrial permeability transition pores (PTP) CYPD, which is required for PTP opening and the mitochondrial membrane permeabilization [102]. A similar effect is triggered by A*β* in the AD brain, which also interacts with CYPD and causes neuronal damage by the mitochondrial membrane permeabilization, loss of the membrane potential, and ATP production [103]. Strikingly, the whole biological process of neurodegeneration can be prevented just by the increase of SIRT3 expression, since it prevents damage of the neuronal mitochondria caused by the co-expression with p53 [101]. The overexpression of p53 can cause 80 % neuronal damage compared to the wild type control, which can be decreased to half by co-expression with SIRT3, while the decline of the neuronal mitochondria membrane potential is also diminished. SIRT3 further protects against the decreasing of cytochrome c activity caused by p53 and prevents neuronal mitochondrial dysfunction [101].

In a conclusion, the multifunctionality in AD includes the regulation of p53 acetylation, which can diminish its negative effect on the mitochondrial membrane potential and prevent neuronal damage. The activators of SIRT3 can effectively decrease the neurodegenerative damages caused by the upregulation of p53, which makes a promise for the future.

### Parkinson’s and Huntington’s disease

Common molecular mechanisms behind PD, HD, and AD and sharing of the substrates were highlighted by the interaction network analysis (Fig. 1A). The involvement of SIRT3 in the PD and HD is, however, supported only by limited data. SIRT3, especially in combination with the physical activity protects against striatal and hippocampal neuron damages in the mouse model of HD [104]. The protein and RNA expression of SIRT3 activated by the exercise increases by about twice compared to the sedentary mouse model of HD [104], which offers options to improve the patient outcomes by the physical activity.

In PD, SIRT3 plays a dominant neuroprotective role in mitochondrial ROS scavenging, which prevents a neurodegenerative change in neurons particularly sensitive to ROS insults. SIRT3 deacetylates and activates SOD2 (cluster IV, Fig. 1A), CYP40 (CYPD), and ATP synthase (cluster II, Fig. 1A). The deacetylation activity mediates the neuroprotection effect of SIRT3, which leads to the quenching of superoxide and efficiently protects against the mitochondrial membrane potential reduction in the PD mouse model [105]. The oxidative damage is also prevented by the regulation of the glutathione peroxidase (GPX) expression by SIRT3 activity [42].

The most utilized PD model is the 1-methyl-4-phenyl-1,2,3,6-tetrahydropyridine (MPTP) mouse model, where the pretreatment causes the inhibition of complex I of MRC creating the condition closely mimicking the condition of Parkinson’s disease. Equally to this model, the complex I MRC activity is also inhibited in SIRT3 KO mice, while the external SIRT3 can restore the activity [48]. SIRT3 knockout further causes the increased dopaminergic neuronal loss in the substantia nigra *pars compacta* (SN) and decreased density of dopaminergic fibers in the striatum signalizing elevated neurodegeneration in comparison to the control MPTP treated mice [42]. The deacetylase controls the permeability of the inner mitochondrial membrane for Ca^2+^, which prevents PTP formation. SIRT3 also increases the cellular ATP production by the prevention of the ATP production arrest, and the neuronal damage through apoptosis [104] [105].

The oxidative damages to the mitochondria occurring during the aging and neurodegenerative changes can be repaired by the removal of the damaged mitochondria through the process of mitophagy. SIRT3 activates mitophagy since the activator metformin stimulates the process of the removal of damaged mitochondria through the SIRT3-PINK1-PARKIN pathway. On the other hand, the SIRT3 inhibitor suppresses the pathway activity [106] [107] and decreases the mitophagy, which demonstrates another neuroprotection activity of SIRT3 activators.

SIRT3 activity mediates ROS scavenging, the regulation of mitophagy and it prevents the mitochondrial membrane potential loss. The neuroprotective effect of the SIRT3 activators in the combination with the active lifestyle is an attractive option for the improvement of cognitive decline during aging and the progressive stages of neurodegeneration.

### Thermogenesis

SIRT3 is importantly involved in the acetylation of the major part of the mitochondrial proteins within the metabolically active brown adipose tissue (BAT), liver, brain, and heart. The highest differential acetylation, however, arises in the wild type versus SIRT3 KO mice in BAT [1].

The SIRT3 substrates involved in clusters I and II (Fig. 1A) are functionally connected to the process of heat production in mammals called thermogenesis, which occurs in BAT. The clusters also overlap with the functions of the complex I, II, and V of MRC (cluster I and cluster II, Fig. 1A) and additional thermogenesis related nodes PPARGC1A, ACSL5, ACSL6, ACTB, and CPT2.

It was previously shown, that SIRT3 knockout does not affect any vital metabolism or the cold resistance of the organism at the standard laboratory maintenance conditions of the experimental animals [1]. During the low-temperature exposure, SIRT3 activates in the brown adipose tissue and the upregulation on the mRNA level further promotes the expression of COX2, COX4, ATP synthase, UCP1, and PPARGC1A. The process leads towards increased mitochondrial respiration and oxygen consumption, which suggests the involvement of SIRT3 in the adaptive thermogenesis [81]. Fasting SIRT3KO mice are extremely cold intolerant and they produce much less hepatic ATP compared to wt mice [19].

Thus, SIRT3 regulates the acetylation of the mitochondrial proteins, while most of the deacetylation occurs in BAT. It contributes to the adaptive thermogenesis of the organisms by the increase of energy production; however, it is not required for the vital metabolism and cold resistance of the animals.

### Non-alcoholic fatty liver disease (NAFLD)

The NAFLD-related SIRT3 substrates are grouped in cluster I of the STRING protein-protein interaction network (Fig. 1A), which correlates with the subunits of the complex I MRC, NADH: ubiquinone oxidoreductase. SIRT3 activation makes a promise for NAFLD therapy, which demonstrates the positive effect of the improved liver mitochondrial energetics in NAFLD patients. The impact is completely abolished in the SIRT3 KO mice, hepatocytes transfected with SIRT3 siRNA, or after the treatment with SIRT3 inhibitor [108]. Another SIRT3 specific drug effective against NAFLD is liraglutide, commonly used for diabetes mellitus type II treatments. It targets SIRT3 and increases its expression, which declines in the high fat (HF) diet-fed mice. The probable liraglutide mode of action is the diminishing of the oxidative stress and activation of the autophagy through the SIRT3-FOXO3a-LC3 pathway [93]. Drug berberine can also reverse the disease symptoms in high-fat diet-induced NAFLD in rats by the activation of the SIRT3-AMPK-ACC (acetyl-CoA carboxylase) pathway [109]. The positive effect of SIRT3 activating drugs for the improvement of NAFLD symptoms is another example of the promising use of SIRT3 as a therapeutical target. The activating of the SIRT3 activity simultaneously in both the brain and liver appear particularly beneficial to counter-balance the negative effect of aging.

## Conclusions

The current substrate interaction network analysis of the main mitochondrial deacetylase SIRT3 highlighted AD, PD, HD, and NAFLD as the diseases the most significantly associated with its deacetylation activity. The most emphasized biological functions of SIRT3 substrates are within the respiratory electron transport chain, TCA cycle, fatty acid, triacylglycerol, and ketone body metabolism.

SIRT3 exhibits several modes of neuroprotective actions in the brain and liver including prevention of mitochondrial damages due to the respiratory electron transfer chain failure, the quenching of ROS, the inhibition of the mitochondrial membrane potential loss, and the regulation of mitophagy. SIRT3 stimulation improves the mitochondrial energetics at the pathological conditions. In the brain tissues, SIRT3 activation performs as a repressor of BACE1, an enzyme that catalyzes the crucial step towards the production of the A*β* peptide, which is considered as the main hallmark of AD. Significantly, SIRT3 also regulates the aggregation of tau through its deacetylation.

The constructed pathway SIRT3-LKB1-AMPK-CREB-PGC-1α-PPARG-BACE1 explains the inhibitory action of SIRT3 on the enzymatic activity of BACE1 and it offers several alternative nodes for the multitargeting by pharmaceuticals. The pathway is closely related to the circadian clock functionality, which also deteriorates during the AD advancement and is the direct causative of the pathological neurodegenerative changes. Additional functions related to AD and the circadian clock also fulfills the pathway SIRT3-FOXO3-CLOCK.

Moreover, SIRT3 also processes p53, protects against p53 associated decrease of cytochrome c activity, and prevents neuronal mitochondrial dysfunction. SIRT3 regulates the metabolic switch which prevents favorable conditions for the cancer cell growth due to the Warburg effect. The majority of SIRT3 deacetylation occurs in BAT, where it contributes to the adaptive thermogenesis of the organisms by the increase of energy production.

The activating of the SIRT3 expression and activity in both the brain and liver seems particularly beneficial to counter-balance the negative effect of aging. The use of the activators in combination with the stimulating effect of regular exercise is an attractive option for the improvement of cognitive decline during aging and the progressive stages of neurodegeneration.

## Acknowledgments

The research was conducted from the financial resources of Biochemworld co., Uppsala County, Sweden.

## Abbreviations

AD: Alzheimer’s disease
CR: calorie restriction
HD: Huntington’s disease
HSC: human stem cell
KO: knockout
MRC: mitochondrial respiratory chain
NAFLD: non-alcoholic fatty liver disease
PD: Parkinson’s disease
PDH: pyruvate dehydrogenase
ROS: reactive oxygen species
TCA: tricarboxylic acid

Tab. 1S The list of the SIRT3 substrates retrieved by the data mining, which were utilized for the pathway enrichment analysis.

Tab. 2S The STRING KEGG pathway enrichment analysis of the SIRT3 substrate interaction network. The analysis was performed using the SIRT3 substrate list as a multiple protein query against the *H. sapiens* database. The settings were selected as follows: active interaction sources – text mining, experiments, and databases; minimum required interaction score – highest confidence (0.9); the maximum number of the interactors to show 1-st and 2-nd shell – none. The terms visualized in Fig. 1A are marked by bold letters.

## References

1. Lombard DB, Alt FW, Cheng H-L, et al. Mammalian Sir2 homolog SIRT3 regulates global mitochondrial lysine acetylation. Mol Cell Biol. 2007; https://doi.org/10.1128/mcb.01636-07

2. McDonnell E, Peterson BS, Bomze HM et al. SIRT3 regulates progression and development of diseases of aging. Trends Endocrinol Metab. 2015; https://doi.org/10.1016/j.tem.2015.06.001

3. Vassilopoulos A, Pennington JD, Andresson T, et al. SIRT3 deacetylates ATP Synthase F1 complex proteins in response to nutrient- and exercise-induced stress. Antioxid Redox Signal. 2013; https://doi.org/10.1089/ars.2013.5420

4. Brown K, Xie S, Qiu X, et al. SIRT3 reverses aging-associated degeneration. Cell Rep. 2013; https://doi.org/10.1016/j.celrep.2013.01.005

5. Braidy N, Poljak A, Grant R, et al. Differential expression of sirtuins in the aging rat brain. Front Cell Neurosci 2015; https://doi.org/10.3389/fncel.2015.00167

6. Weir HJM, Murray TK, Kehoe PG, et al. CNS SIRT3 expression is altered by reactive oxygen species and in Alzheimer’s disease. PLoS One 2012; https://doi.org/10.1371/journal.pone.0048225

7. Oti M. Predicting disease genes using protein-protein interactions. J Med Genet. 2006;. https://doi.org/10.1136/jmg.2006.041376

8. Poulose N, Raju R. Sirtuin regulation in aging and injury. Biochim Biophys Acta - Mol Basis Dis. 2015; https://doi.org/10.1016/j.bbadis.2015.08.017

9. Hallows WC, Lee S, Denu JM. Sirtuins deacetylate and activate mammalian acetyl-CoA synthetases. Proc Natl Acad Sci U S A. 2006; https://doi.org/10.1073/pnas.0604392103

10. Shimazu T, Hirschey MD, Hua L, et al. SIRT3 deacetylates mitochondrial 3-hydroxy-3-methylglutaryl CoA synthase 2 and regulates ketone body production. Cell Metab. 2010; https://doi.org/10.1016/j.cmet.2010.11.003

11. Bharathi SS, Zhang Y, Mohsen AW, et al. Sirtuin 3 (SIRT3) protein regulates long-chain acyl-CoA dehydrogenase by deacetylating conserved lysines near the active site. J Biol Chem. 2013; https://doi.org/10.1074/jbc.M113.510354

12. Finley LWSS, Carracedo A, Lee J, et al. SIRT3 opposes reprogramming of cancer cell metabolism through HIF1α destabilization. Cancer Cell. 2011; https://doi.org/10.1016/j.ccr.2011.02.014

13. Sundaresan NR, Samant SA, Pillai VB, et al. SIRT3 is a stress-responsive deacetylase in cardiomyocytes that protects cells from stress-mediated cell death by deacetylation of Ku70. Mol Cell Biol. 2008; https://doi.org/10.1128/mcb.00426-08

14. Qiu X, Brown K, Hirschey MD, et al. Calorie restriction reduces oxidative stress by SIRT3-mediated SOD2 activation. Cell Metab. 2010; https://doi.org/10.1016/j.cmet.2010.11.015

15. Yu W, Dittenhafer-Reed KE, Denu JM. SIRT3 protein deacetylates isocitrate dehydrogenase 2 (IDH2) and regulates mitochondrial redox status. J Biol Chem. 2012; https://doi.org/10.1074/jbc.M112.355206

16. Schlicker C, Gertz M, Papatheodorou P, et al. Substrates and regulation mechanisms for the human mitochondrial sirtuins Sirt3 and Sirt5. J Mol Biol. 2008; https://doi.org/10.1016/j.jmb.2008.07.048

17. Pillai VB, Sundaresan NR, Kim G, et al. Exogenous NAD blocks cardiac hypertrophic response via activation of the SIRT3-LKB1-AMP-activated kinase pathway. J Biol Chem. 2010. https://doi.org/10.1074/jbc.M109.077271

18. Yang Y, Cimen H, Han MJ, et al. NAD+-dependent deacetylase SIRT3 regulates mitochondrial protein synthesis by deacetylation of the ribosomal protein MRPL10. J Biol Chem. 2010;.https://doi.org/10.1074/jbc.M109.053421

19. Hirschey MD, Shimazu T, Goetzman E, et al. SIRT3 regulates mitochondrial fatty-acid oxidation by reversible enzyme deacetylation. Nature 2010; https://doi.org/10.1038/nature08778

20. Vassilopoulos A, Pennington JD, Andresson T, et al. SIRT3 deacetylates ATP synthase F1 complex proteins in response to nutrient- and exercise-induced stress. Antioxid Redox Signal. 2014; https://doi.org/10.1089/ars.2013.5420

21. Xue L, Xu F, Meng L, et al. Acetylation-dependent regulation of mitochondrial ALDH2 activation by SIRT3 mediates acute ethanol-induced eNOS activation. FEBS Lett. 2011; https://doi.org/10.1016/j.febslet.2011.11.031

22. Wang Z, Inuzuka H, Zhong J, et al. Identification of acetylation-dependent regulatory mechanisms that govern the oncogenic functions of Skp2. Oncotarget. 2012; https://doi.org/10.18632/oncotarget.740

23. Tseng AHH, Shieh SS, Wang DL SIRT3 deacetylates FOXO3 to protect mitochondria against oxidative damage. Free Radic Biol Med. 2013; https://doi.org/10.1016/j.freeradbiomed.2013.05.002

24. Jing E, O’Neill BT, Rardin MJ, et al. Sirt3 regulates metabolic flexibility of skeletal muscle through reversible enzymatic deacetylation. Diabetes. 2013; https://doi.org/10.2337/db12-1650

25. Cheng Y, Ren X, Gowda ASP, et al. Interaction of Sirt3 with OGG1 contributes to repair of mitochondrial DNA and protects from apoptotic cell death under oxidative stress. Cell Death Dis. 2013; https://doi.org/10.1038/cddis.2013.254

26. Samant SA, Zhang HJ, Hong Z, et al. SIRT3 Deacetylates and Activates OPA1 To Regulate Mitochondrial Dynamics during Stress. Mol Cell Biol. 2014; https://doi.org/10.1128/mcb.01483-13

27. Lu Z, Chen Y, Aponte AM, et al. Prolonged fasting identifies heat shock protein 10 as a sirtuin 3 substrate: Elucidating a new mechanism linking mitochondrial protein acetylation to fatty acid oxidation enzyme folding and function. J Biol Chem. 2015; https://doi.org/10.1074/jbc.M114.606228

28. Rauh D, Fischer F, Gertz M, et al. An acetylome peptide microarray reveals specificities and deacetylation substrates for all human sirtuin isoforms. Nat Commun. 2013; https://doi.org/10.1038/ncomms3327

29. Yang H, Zhou L, Shi Q, et al. SIRT 3-dependent GOT 2 acetylation status affects the malate–aspartate NADH shuttle activity and pancreatic tumor growth. EMBO J. 2015; https://doi.org/10.15252/embj.201591041

30. Rardin MJ, Newman JC, Held JM, et al. Label-free quantitative proteomics of the lysine acetylome in mitochondria identifies substrates of SIRT3 in metabolic pathways. Proc Natl Acad Sci U S A. 2013; https://doi.org/10.1073/pnas.1302961110

31. Hebert AS, Dittenhafer-Reed KE, Yu W, et al. Calorie restriction and SIRT3 trigger global reprogramming of the mitochondrial protein acetylome. Mol Cell. 2012; https://doi.org/10.1016/j.molcel.2012.10.024

32. Sol EM, Wagner SA, Weinert BT, et al. Proteomic investigations of lysine acetylation identify diverse substrates of mitochondrial deacetylase Sirt3. PLoS One. 2012; https://doi.org/10.1371/journal.pone.0050545

33. Zuberi K, Franz M, Rodriguez H, et al. GeneMANIA prediction server 2013 update. Nucleic Acids Res. 2013; https://doi.org/10.1093/nar/gkt533

34. Warde-Farley D, Donaldson SL, Comes O, et al. The GeneMANIA prediction server: Biological network integration for gene prioritization and predicting gene function. Nucleic Acids Res. 2010; https://doi.org/10.1093/nar/gkq537

35. Franz M, Rodriguez H, Lopes C, et al. GeneMANIA update 2018. Nucleic Acids Res. 2018; https://doi.org/10.1093/nar/gky311

36. Lopes CT, Franz M, Kazi F, et al. Cytoscape Web: an interactive web-based network browser. Bioinformatics. 2010; https://doi.org/10.1093/bioinformatics/btq430

37. Szklarczyk D, Gable AL, Lyon D, et al. STRING v11: Protein-protein association networks with increased coverage, supporting functional discovery in genome-wide experimental datasets. Nucleic Acids Res. 2019; https://doi.org/10.1093/nar/gky1131

38. Enright AJ, Van Dongen S, Ouzounis CA. An efficient algorithm for large-scale detection of protein families. Nucleic Acids Res. 2020; https://doi.org/10.1093/nar/30.7.1575

39. Chin C-HH, Chen S-HH, Wu H-HH, et al. CytoHubba: identifying hub objects and sub-networks from complex interactome. BMC Syst Biol 8 Suppl. 2014; https://doi.org/10.1186/1752-0509-8-S4-S11

40. Bader GD, Hogue CWV. An automated method for finding molecular complexes in large protein interaction networks. BMC Bioinformatics. 2003; https://doi.org/10.1186/1471-2105-4-2

41. Kutmon M, Lotia S, Evelo CT, Pico AR. WikiPathways App for Cytoscape: Making biological pathways amenable to network analysis and visualization. F1000Research. 2014; https://doi.org/10.12688/f1000research.4254.2

42. Liu L, Peritore C, Ginsberg J, et al. SIRT3 Attenuates MPTP-induced nigrostriatal degeneration via enhancing mitochondrial antioxidant capacity. Neurochem Res. 2015. https://doi.org/10.1007/s11064-014-1507-8

43. Franceschini A, Szklarczyk D, Frankild S, et al. STRING v9.1: Protein-protein interaction networks, with increased coverage and integration. Nucleic Acids Res. 2013; https://doi.org/10.1093/nar/gks1094

44. Szklarczyk D, Morris JH, Cook H, et al. The STRING database in 2017: quality-controlled protein-protein association networks, made broadly accessible. Nucleic Acids Res. 2017; https://doi.org/10.1093/nar/gkw937

45. Bader JS, Chaudhuri A, Rothberg JM et al. Gaining confidence in high-throughput protein interaction networks. Nat Biotechnol. 2004; https://doi.org/10.1038/nbt924

46. Dongen SM van. Graph clustering by flow simulation. Ph.D. thesis, Univ Utr. 2000. https://dspace.library.uu.nl/bitstream/handle/1874/848/full.pdf?sequence=1&isAllowed=y; Accessed October 2, 2020.

47. Manczak M, Park BS, Jung Y et al. Differential Expression of Oxidative Phosphorylation Genes in Patients With Alzheimer’s Disease: Implications for Early Mitochondrial Dysfunction and Oxidative Damage. Neuromolecular Med. 2004; https://doi.org/10.1385/NMM:5:2:147

48. Ahn BH, Kim HS, Song S, et al. A role for the mitochondrial deacetylase Sirt3 in regulating energy homeostasis. Proc Natl Acad Sci U S AA. 2008; https://doi.org/10.1073/pnas.0803790105

49. Bao J, Scott I, Lu Z, et al. SIRT3 is regulated by nutrient excess and modulates hepatic susceptibility to lipotoxicity. Free Radic Biol Med. 2010; https://doi.org/10.1016/j.freeradbiomed.2010.07.009

50. Cimen H, Han M-J, Yang Y, et al. Regulation of succinate dehydrogenase activity by SIRT3 in mammalian mitochondria. Biochemistry. 2010; https://doi.org/10.1021/bi901627u

51. Finley LWS, Haas W, Desquiret-Dumas V, et al. Succinate dehydrogenase is a direct target of sirtuin 3 deacetylase activity. PLoS One. 2011; https://doi.org/10.1371/journal.pone.0023295

52. Bellizzi D, Rose G, Cavalcante P, et al. A novel VNTR enhancer within the SIRT3 gene, a human homologue of SIR2, is associated with survival at oldest ages. Genomics. 2005; https://doi.org/10.1016/j.ygeno.2004.11.003

53. Yang W, Nagasawa K, Münch C, et al. Mitochondrial Sirtuin Network Reveals Dynamic SIRT3-Dependent Deacetylation in Response to Membrane Depolarization. Cell. 2016; https://doi.org/10.1016/j.cell.2016.10.016

54. Bubber P, Haroutunian V, Fisch G, et al. Mitochondrial abnormalities in Alzheimer brain: Mechanistic implications. Ann Neurol. 2005;https://doi.org/10.1002/ana.20474

55. Ozden O, Park S-HH, Wagner BA, et al. SIRT3 deacetylates and increases pyruvate dehydrogenase activity in cancer cells. Free Radic Biol Med. 2014; https://doi.org/10.1016/j.freeradbiomed.2014.08.001

56. Warburg O, Wind F, Negelein E. The metabolism of tumors in the body. J Gen Physiol. 1927; https://doi.org/10.1085/jgp.8.6.519

57. Ozden O, Park S-H, Wagner BA, et al. SIRT3 deacetylates and increases pyruvate dehydrogenase activity in cancer cells. Free Radic Biol Med. 2014; https://doi.org/10.1016/j.freeradbiomed.2014.08.001

58. Stacpoole PW. The pyruvate dehydrogenase complex as a therapeutic target for age-related diseases. Aging Cell. 2012; https://doi.org/10.1111/j.1474-9726.2012.00805.x

59. Zhou Q, Lam PY, Han D et al. Activation of c-Jun-N-terminal kinase and decline of mitochondrial pyruvate dehydrogenase activity during brain aging. FEBS Lett. 2009; https://doi.org/10.1016/j.febslet.2009.02.043

60. Scher MB, Vaquero A, Reinberg D. SirT3 is a nuclear NAD+-dependent histone deacetylase that translocates to the mitochondria upon cellular stress. Genes Dev. 2007; https://doi.org/10.1101/gad.1527307

61. Cooper HM, Spelbrink JN. The human SIRT3 protein deacetylase is exclusively mitochondrial. Biochem J. 2008; https://doi.org/10.1042/BJ20071624

62. Gurd BJ, Holloway GP, Yoshida Y et al. In mammalian muscle, SIRT3 is present in mitochondria and not in the nucleus; and SIRT3 is upregulated by chronic muscle contraction in an adenosine monophosphate-activated protein kinase–independent manner. Metabolism, 2012; https://doi.org/10.1016/j.metabol.2011.09.016

63. Zhang X, Cao R, Niu J, et al. Molecular basis for hierarchical histone de-β-hydroxybutyrylation by SIRT3. Cell Discov. 2019; https://doi.org/10.1038/s41421-019-0103-0

64. Sengupta A, Haldar D. Human sirtuin 3 (SIRT3) deacetylates histone H3 lysine 56 to promote nonhomologous end-joining repair. DNA Repair (Amst). 2018; https://doi.org/https://doi.org/10.1016/j.dnarep.2017.11.003

65. Vedrenne V, Gowher A, De Lonlay P, et al. Mutation in PNPT1, which encodes a polyribonucleotide nucleotidyltransferase, impairs RNA import into mitochondria and causes respiratory-chain deficiency. Am J Hum Genet. 2012; https://doi.org/10.1016/j.ajhg.2012.09.001

66. Palmieri L, Pardo B, Lasorsa FM, et al. Citrin and aralar1 are Ca2+-stimulated aspartate/glutamate transporters in mitochondria. EMBO J. 2001; https://doi.org/10.1093/emboj/20.18.5060

67. Galmiche L, Serre V, Beinat MA. Toward genotype phenotype correlations in GFM1 mutations. Mitochondrion. 2012; https://doi.org/10.1016/j.mito.2011.09.007

68. Fukumura S, Ohba C, Watanabe T, et al. Compound heterozygous GFM2 mutations with Leigh syndrome complicated by arthrogryposis multiplex congenita. J Hum Genet. 2015; https://doi.org/10.1038/jhg.2015.57

69. Perli E, Pisano A, Glasgow RIC, et al. Novel compound mutations in the mitochondrial translation elongation factor (TSFM) gene cause severe cardiomyopathy with myocardial fibro-adipose replacement. Sci Rep. 2019; https://doi.org/10.1038/s41598-019-41483-9

70. Shi H, Hayes M, Kirana C, Miller R, Keating J, Macartney-Coxson D, Stubbs R. TUFM is a potential new prognostic indicator for colorectal carcinoma. Pathology. 2012; https://doi:10.1097/PAT.0b013e3283559cbe.

71. Schrader M, Kamoshita M, Islinger M. Organelle interplay-peroxisome interactions in health and disease. J Inherit Metab Dis. 2020; https://doi.org/10.1002/jimd.12083

72. Hirschey MD, Shimazu T, Jing E, et al. SIRT3 deficiency and mitochondrial protein hyperacetylation accelerate the development of the metabolic syndrome. Mol Cell. 2011; https://doi.org/10.1016/j.molcel.2011.07.019

73. Yechoor VK, Patti ME, Ueki K, et al. Distinct pathways of insulin-regulated versus diabetes-regulated gene expression: An in vivo analysis in MIRKO mice. Proc Natl Acad Sci U S A. 2004; https://doi.org/10.1073/pnas.0407574101

74. Tyagi A, Nguyen CU, Chong T, et al. SIRT3 deficiency-induced mitochondrial dysfunction and inflammasome formation in the brain. Sci Rep. 2008; https://doi.org/10.1038/s41598-018-35890-7

75. Xu WL, Atti AR, Gatz M, et al. Midlife overweight and obesity increase late-life dementia risk. Neurology. 2011; https://doi.org/10.1212/WNL.0b013e3182190d09

76. Study MA, Spauwen PJJ, Köhler S, et al. Effects of Type 2 Diabetes on 12-Year Cognitive Change. Diabetes Care. 2013; https://doi.org/10.2337/dc12-0746

77. Pillai VB, Sundaresan NR, Kim G, et al. Exogenous NAD blocks cardiac hypertrophic response via activation of the SIRT3-LKB1-AMP-activated kinase pathway. J Biol Chem. 2010; https://doi.org/10.1074/jbc.M109.077271

78. Palacios OM, Carmona JJ, Michan S, et al. Diet and exercise signals regulate SIRT3 and activate AMPK and PGC-1alpha in skeletal muscle. Aging (Albany NY). 2009; https://doi.org/10.18632/aging.100075

79. Ramesh S, Govindarajulu M, Lynd T, et al. SIRT3 activator Honokiol attenuates β-Amyloid by modulating amyloidogenic pathway. PLoS One. 2018; https://doi.org/10.1371/journal.pone.0190350

80. Katsouri L, Parr C, Bogdanovic N, et al. PPARγ co-activator-1α (PGC-1α) reduces amyloid-β generation through a PPARγ-dependent mechanism. J Alzheimer’s Dis. 2011; https://doi.org/10.3233/JAD-2011-101356

81. Shi T, Wang F, Stieren E, Tong Q. SIRT3, a mitochondrial sirtuin deacetylase, regulates mitochondrial function and thermogenesis in brown adipocytes. J Biol Chem. 2005; https://doi.org/10.1074/jbc.M414670200

82. Wolfe DM, Lee J, Kumar A, et al. Autophagy failure in Alzheimer’s disease and the role of defective lysosomal acidification. Eur J Neurosci. 2013; https://doi.org/10.1111/ejn.12169

83. Li S, Dou X, Ning H, et al. Sirtuin 3 acts as a negative regulator of autophagy dictating hepatocyte susceptibility to lipotoxicity. Hepatology. 2017; https://doi.org/10.1002/hep.29229

84. Qin W, Haroutunian V, Katsel P, et al. PGC-1α expression decreases in the Alzheimer disease brain as a function of dementia. Arch Neurol. 2009; https://doi.org/10.1001/archneurol.2008.588

85. Wu Z, Puigserver P, Andersson U, et al. Mechanisms controlling mitochondrial biogenesis and respiration through the thermogenic coactivator PGC-1. Cell. 1998; https://doi.org/10.1109/freq.2001.956373

86. Liu C, Li S, Liu T, et al. Transcriptional coactivator PGC-1α integrates the mammalian clock and energy metabolism. Nature. 2007; https://doi.org/10.1038/nature05767

87. Song C, Zhao J, Zhang J, et al. SIRT3-dependent mitochondrial oxidative stress in sodium fluoride-induced hepatotoxicity and salvage by melatonin. BioRxiv. 2017; https://doi.org/10.1101/107813

88. Cermakian N, Waddington Lamont E, Boudreau P et al. Circadian Clock Gene Expression in Brain Regions of Alzheimer’s Disease Patients and Control Subjects. J Biol Rhythms. 2011; https://doi.org/10.1177/0748730410395732

89. Kondratova AA, Kondratov RV. The circadian clock and pathology of the ageing brain. Nat Rev Neurosci. 2012; https://doi.org/10.1038/nrn3208

90. Mauvoisin D, Atger F, Dayon L, et al. Circadian and Feeding Rhythms Orchestrate the Diurnal Liver Acetylome. Cell Rep. 2017; 1729–1743. https://doi.org/https://doi.org/10.1016/j.celrep.2017.07.065

91. Tomita T. Aberrant proteolytic processing and therapeutic strategies in Alzheimer disease. Adv Biol Regul. 2017; https://doi.org/10.1016/j.jbior.2017.01.001

92. Chaves I, van der Horst GTJ, Schellevis R, et al. Insulin-FOXO3 signaling modulates circadian rhythms via regulation of clock transcription. Curr Biol. 2014; https://doi.org/10.1016/j.cub.2014.04.018

93. Tong W, Ju L, Qiu M, et al. Liraglutide ameliorates non-alcoholic fatty liver disease by enhancing mitochondrial architecture and promoting autophagy through the SIRT1/SIRT3–FOXO3a pathway. Hepatol Res. 2016; https://doi.org/10.1111/hepr.12634

94. Jacobs KM, Pennington JD, Bisht KS, et al. SIRT3 interacts with the daf-16 homolog FOXO3a in the Mitochondria, as well as increases FOXO3a Dependent Gene expression. Int J Biol Sci. 2008; https://doi.org/10.7150/ijbs.4.291

95. Li H, Jia J, Wang W, et al. Honokiol alleviates cognitive deficits of Alzheimer’s disease (PS1 V97L) transgenic mice by activating mitochondrial SIRT3. J Alzheimer’s Dis. 2008; https://doi.org/10.3233/JAD-180126

96. Cheng A, Wang J, Ghena N, et al. SIRT3 Haploinsufficiency Aggravates Loss of GABAergic Interneurons and Neuronal Network Hyperexcitability in an Alzheimer’s Disease Model. J Neurosci. 2020; https://doi.org/10.1523/JNEUROSCI.1446-19.2019

97. Tracy TE, Sohn PD, Minami SS, et al. Acetylated tau obstructs KIBRA-mediated signaling in synaptic plasticity and promotes tauopathy-related memory loss. Neuron. 2016; https://doi.org/https://doi.org/10.1016/j.neuron.2016.03.005

98. Cohen TJ, Guo JL, Hurtado DE, et al. The acetylation of tau inhibits its function and promotes pathological tau aggregation. Nat Commun. 2011; https://doi.org/10.1038/ncomms1255

99. Yin J, Han P, Song M, et al. Amyloid-β Increases Tau by Mediating Sirtuin 3 in Alzheimer’s Disease. Mol Neurobiol. 2018; https://doi.org/10.1007/s12035-018-0977-0

100. Li, S., Yin, J., Nielsen, M. et al. Sirtuin 3 mediates tau deacetylation. J. Alzheimer’s Dis. 2019; https://doi.org/10.3233/JAD-190014

101. Lee J, Kim Y, Liu T, et al. SIRT3 deregulation is linked to mitochondrial dysfunction in Alzheimer’s disease. Aging Cell. 2018; 17:1–12. https://doi.org/10.1111/acel.12679

102. Vaseva A V., Marchenko ND, Ji K, et al. P53 opens the mitochondrial permeability transition pore to trigger necrosis. Cell. 2012; https://doi.org/10.1016/j.cell.2012.05.014

103. Du H, Guo L, Fang F, et al. Cyclophilin D deficiency attenuates mitochondrial and neuronal perturbation and ameliorates learning and memory in Alzheimer’s disease. 2009; https://doi.org/10.1038/nm.1868.Cyclophilin

104. Cheng A, Yang Y, Zhou Y, et al. Mitochondrial SIRT3 Mediates Adaptive Responses of Neurons to Exercise and Metabolic and Excitatory Challenges. Cell Metab. 2016; https://doi.org/10.1016/j.cmet.2015.10.013

105. Zhang X, Ren X, Zhang Q, et al. PGC-1α/ERRα-Sirt3 pathway regulates DAergic neuronal death by directly deacetylating SOD2 and ATP synthase β. Antioxidants Redox Signal. 2016; https://doi.org/10.1089/ars.2015.6403

106. Wang C, Yang Y, Zhang Y, et al. Protective effects of metformin against osteoarthritis through upregulation of SIRT3-mediated PINK1/Parkin-dependent mitophagy in primary chondrocytes. Biosci Trends. 2018; https://doi.org/10.5582/bst.2018.01263

107. Yu W, Gao B, Li N, et al. Sirt3 deficiency exacerbates diabetic cardiac dysfunction: Role of Foxo3A-Parkin-mediated mitophagy. Biochim Biophys Acta - Mol Basis Dis. 2017; https://doi.org/10.1016/j.bbadis.2016.10.021

108. Zeng X, Yang J, Hu O, et al. Dihydromyricetin ameliorates nonalcoholic fatty liver disease by improving mitochondrial respiratory capacity and redox homeostasis through modulation of SIRT3 signaling. Antioxid Redox Signal 2018; https://doi.org/10.1089/ars.2017.7172

109. Zhang Y, Deng Y, Tang K, et al. Berberine ameliorates high-fat diet-induced nonalcoholic fatty liver disease in rats via activation of SIRT3/AMPK/ACC Pathway. Curr Med Sci. 2019; https://doi.org/10.1007/s11596-019-1997-3

