## Supplementary material for "The function of SIRT3 explored through the substrate interaction network": Tab. 2S

| Term description | FDR | Matching proteins in the network |
|---|---|---|
| Metabolic pathways | 2.40e-81 | AADAT,AASS,ABAT,ACAA2, ACADL, ACADM, ACADS, ACADSB, ACADVL, ACAT1, ACO2, ACOT1, ACOT2, ACOX1, ACSL5, ACSL6, ACSM1, ACSM3, ACSM5, ACSS2, ACSS3, AGMAT, AGXT2, AK4, ALAS1, ALDH1B1, ALDH2, ALDH4A1, ALDH5A1, ALDH6A1, ALDOB, AMACR, AMT, APRT, ARG1, ATP5A1, ATP5B, ATP5C1, ATP5E, ATP5F1, ATP5H, ATP5J, ATP5L, ATP5O, AUH, BCKDHA, BDH1, CHDH, COX4I1, COX7C, CPS1, CS, CYCS, CYP1A2, CYP27A1, DBT, DHRS4, DLAT, DLD, DLST, DMGDH, ECHS1, EHHADH, ENO1, EPHX2, FAHD1, FASN, FDPS, FECH, FH, GBE1, GCDH, GLDC, GLS, GLS2, GLUD1, GOT2, GPAM, GPI, GPT2, H6PD, HADH, HADHA, HADHB, HIBADH, HIBCH, HMGCL, HMGCS2, HSD17B10, HSD17B12, IDH1, IDH2, IDH3A, IDH3G, IVD, MCCC1, MCCC2, MCEE, MDH2, ME1, ME3, MECR, MLYCD, MMAB, MPST, MT-ATP8, MTHFD1L, MUT, NDUFA10, NDUFA12, NDUFA13, NDUFA2, NDUFA5, NDUFA7, NDUFA9, NDUFAB1, NDUFB11, NDUFB3, NDUFS1, NDUFS2, NDUFS3, NDUFS6, NDUFS7, NDUFS8, NDUFV1, NDUFV2, NDUFV3, NNT, OAT, OGDH, OTC, OXSM, PC, PCCA, PCCB, PDHA1, PDHB, PDHX, PRODH2, PYCR2, PYGL, SARDH, SDHA, SDHB, SHMT2, SLC27A5, SUCLA2, SUCLG1, SUCLG2, TST, UGP2, UQCRC2, UQCRFS1, UQCRQ |
| Carbon metabolism | 4.12e-43 | ACADM, ACADS, ACAT1, ACO2, ACSS2, ALDH6A1, ALDOB, AMT, CAT, CPS1, CS, DLAT, DLD, DLST, ECHS1, EHHADH, ENO1, FH, GLDC, GLUD1, GOT2, GPI, GPT2, H6PD, HADHA, HIBCH, IDH1, IDH2, IDH3A, IDH3G, MCEE, MDH2, ME1, ME2, ME3, MUT, OGDH, PC, PCCA, PCCB, PDHA1, PDHB, SDHA, SDHB, SHMT2, SUCLA2, SUCLG1, SUCLG2 |
| **Valine, leucine, and isoleucine degradation** | 1.44e-33 | ABAT, ACAA2, ACADM, ACADS, ACADSB, ACAT1, ACSF3, AGXT2, ALDH1B1, ALDH2, ALDH6A1, AUH, BCKDHA, DBT, DLD, ECHS1, EHHADH, HADH, HADHA, |

| | | |
|---|---|---|
| | | HADHB, HIBADH, HIBCH, HMGCL, HMGCS2, HSD17B10, IVD, MCCC1, MCCC2, MCEE, MUT, PCCA, PCCB |
| **Parkinson's disease** | 7.21e-33 | ATP5A1, ATP5B, ATP5C1, ATP5E, ATP5F1, ATP5H, ATP5J, ATP5O, COX4I1, COX7A2, COX7C, CYCS, MT-ATP8, NDUFA10, NDUFA12, NDUFA13, NDUFA2, NDUFA5, NDUFA7, NDUFA9, NDUFAB1, NDUFB11, NDUFB3, NDUFS1, NDUFS2, NDUFS3, NDUFS6, NDUFS7, NDUFS8, NDUFV1, NDUFV2, NDUFV3, PARK7, PPIF, SDHA, SDHB, SLC25A4, SLC25A5, SLC25A6, UQCRC2, UQCRFS1, UQCRQ |
| **Huntington's disease** | 1.49e-31 | ATP5A1, ATP5B, ATP5C1, ATP5E, ATP5F1, ATP5H, ATP5J, ATP5O, COX4I1, COX7A2, COX7C, CYCS, MT-ATP8, NDUFA10, NDUFA12, NDUFA13, NDUFA2, NDUFA5, NDUFA7, NDUFA9, NDUFAB1, NDUFB11, NDUFB3, NDUFS1, NDUFS2, NDUFS3, NDUFS6, NDUFS7, NDUFS8, NDUFV1, NDUFV2, NDUFV3, PPARGC1A, PPIF, SDHA, SDHB, SLC25A4, SLC25A5, SLC25A6, SOD2, TFAM, TP53, UQCRC2, UQCRFS1, UQCRQ |
| **Oxidative phosphorylation** | 1.48e-29 | ATP5A1, ATP5B, ATP5C1, ATP5E, ATP5F1, ATP5H, ATP5J, ATP5L, ATP5O, COX4I1, COX7A2, COX7C, MT-ATP8, NDUFA10, NDUFA12, NDUFA13, NDUFA2, NDUFA5, NDUFA7, NDUFA9, NDUFAB1, NDUFB11, NDUFB3, NDUFS1, NDUFS2, NDUFS3, NDUFS6, NDUFS7, NDUFS8, NDUFV1, NDUFV2, NDUFV3, PPA2, SDHA, SDHB, UQCRC2, UQCRFS1, UQCRQ |
| **Thermogenesis** | 5.47e-29 | ACSL5, ACSL6, ACTB, ATP5A1, ATP5B, ATP5C1, ATP5E, ATP5F1, ATP5H, ATP5J, ATP5L, ATP5O, COX4I1, COX7A2, COX7C, CPT2, MT-ATP8, NDUFA10, NDUFA12, NDUFA13, NDUFA2, NDUFA5, NDUFA7, NDUFA9, NDUFAB1, NDUFAF2, NDUFAF4, NDUFB11, NDUFB3, NDUFS1, NDUFS2, NDUFS3, NDUFS6, NDUFS7, NDUFS8, NDUFV1, NDUFV2, NDUFV3, PPARGC1A, SDHA, SDHB, SLC25A20, UQCRC2, UQCRFS1, UQCRQ |
| **Alzheimer's disease** | 2.73e-26 | ATP5A1, ATP5B, ATP5C1, ATP5E, ATP5F1, ATP5H, ATP5J, ATP5O, COX4I1, COX7A2, COX7C, CYCS, HSD17B10, MT-ATP8, NDUFA10, NDUFA12, NDUFA13, NDUFA2, NDUFA5, NDUFA7, NDUFA9, NDUFAB1, NDUFB11, NDUFB3, NDUFS1, NDUFS2, |

| | | |
|---|---|---|
| | | NDUFS3, NDUFS6, NDUFS7, NDUFS8, NDUFV1, NDUFV2, NDUFV3, SDHA, SDHB, UQCRC2, UQCRFS1, UQCRQ |
| **Propanoate metabolism** | 1.54e-23 | ABAT, ACADM, ACAT1, ACSS2, ACSS3, ALDH6A1, BCKDHA, DBT, DLD, ECHDC1, ECHS1, EHHADH, HADHA, HIBCH, MCEE, MLYCD, MUT, PCCA, PCCB, SUCLA2, SUCLG1, SUCLG2 |
| **Citrate cycle (TCA cycle)** | 3.09e-21 | ACO2, CS, DLAT, DLD, DLST, FH, IDH1, IDH2, IDH3A, IDH3G, MDH2, OGDH, PC, PDHA1, PDHB, SDHA, SDHB, SUCLA2, SUCLG1, SUCLG2 |
| **Fatty acid degradation** | 4.84e-20 | ACAA2, ACADL, ACADM, ACADS, ACADSB, ACADVL, ACAT1, ACOX1, ACSL5, ACSL6, ALDH1B1, ALDH2, CPT2, ECHS1, ECI1, ECI2, EHHADH, GCDH, HADH, HADHA, HADHB |
| Fatty acid metabolism | 3.36e-18 | ACAA2, ACADL, ACADM, ACADS, ACADSB, ACADVL, ACAT1, ACOX1, ACSL5, ACSL6, CPT2, ECHS1, EHHADH, FASN, HADH, HADHA, HADHB, HSD17B12, MECR, OXSM |
| **Non-alcoholic fatty liver disease (NAFLD)** | 1.30e-17 | COX4I1, COX7A2, COX7C, CYCS, NDUFA10, NDUFA12, NDUFA13, NDUFA2, NDUFA5, NDUFA7, NDUFA9, NDUFAB1, NDUFB11, NDUFB3, NDUFS1, NDUFS2, NDUFS3, NDUFS6, NDUFS7, NDUFS8, NDUFV1, NDUFV2, NDUFV3, SDHA, SDHB, UQCRC2, UQCRFS1, UQCRQ |
| Butanoate metabolism | 1.10e-13 | ABAT, ACADS, ACAT1, ACSM1, ACSM3, ACSM5, ALDH5A1, BDH1, ECHS1, EHHADH, HADH, HADHA, HMGCL, HMGCS2 |
| Pyruvate metabolism | 2.39e-13 | ACAT1, ACSS2, ALDH1B1, ALDH2, DLAT, DLD, FH, LDHD, MDH2, ME1, ME2, ME3, PC, PDHA1, PDHB |
| Glyoxylate and dicarboxylate metabolism | 1.87e-12 | ACAT1, ACO2, AMT, CAT, CS, DLD, GLDC, MCEE, MDH2, MUT, PCCA, PCCB, SHMT2 |
| Peroxisome | 1.63e-11 | ACOX1, ACSL5, ACSL6, AMACR, CAT, DHRS4, ECH1, ECI2, EHHADH, EPHX2, HMGCL, IDH1, IDH2, MLYCD, PEX2, PRDX5, SOD2 |
| Biosynthesis of amino acids | 3.33e-11 | ACO2, ALDOB, ARG1, CPS1, CS, ENO1, GOT2, GPT2, IDH1, IDH2, IDH3A, IDH3G, OTC, PC, PYCR2, SHMT2 |

| | | |
|---|---|---|
| Tryptophan metabolism | 8.93e-10 | AADAT, ACAT1, ALDH1B1, ALDH2, CAT, CYP1A2, ECHS1, EHHADH, GCDH, HADH, HADHA, OGDH |
| Retrograde endocannabinoid signaling | 1.42e-09 | NDUFA10, NDUFA12, NDUFA13, NDUFA2, NDUFA5, NDUFA7, NDUFA9, NDUFAB1, NDUFB11, NDUFB3, NDUFS1, NDUFS2, NDUFS3, NDUFS6, NDUFS7, NDUFS8, NDUFV1, NDUFV2, NDUFV3 |
| 2-Oxocarboxylic acid metabolism | 3.46e-09 | AADAT, ACO2, CS, GOT2, GPT2, IDH1, IDH2, IDH3A, IDH3G |
| Glycine, serine, and threonine metabolism | 8.28e-09 | AGXT2, ALAS1, AMT, CHDH, DLD, DMGDH, GCAT, GLDC, GNMT, SARDH, SHMT2 |
| beta-Alanine metabolism | 1.57e-08 | ABAT, ACADM, ALDH1B1, ALDH2, ALDH6A1, ECHS1, EHHADH, HADHA, HIBCH, MLYCD |
| Lysine degradation | 3.24e-08 | AADAT, AASS, ACAT1, ALDH1B1, ALDH2, DLST, ECHS1, EHHADH, GCDH, HADH, HADHA, OGDH |
| Alanine, aspartate, and glutamate metabolism | 3.86e-08 | ABAT, AGXT2, ALDH4A1, ALDH5A1, CPS1, GLS, GLS2, GLUD1, GOT2, GPT2 |
| Fatty acid elongation | 4.15e-08 | ACAA2, ACOT1, ACOT2, ECHS1, HADH, HADHA, HADHB, HSD17B12, MECR |
| Arginine biosynthesis | 1.48e-07 | ARG1, CPS1, GLS, GLS2, GLUD1, GOT2, GPT2, OTC |
| PPAR signaling pathway | 2.00e-07 | ACADL, ACADM, ACOX1, ACSL5, ACSL6, CPT2, CYP27A1, DBI, EHHADH, HMGCS2, ME1, SLC27A5 |
| Arginine and proline metabolism | 4.45e-06 | AGMAT, ALDH1B1, ALDH2, ALDH4A1, ARG1, GOT2, OAT, PRODH2, PYCR2 |
| Glycolysis / Gluconeogenesis | 7.55e-06 | ACSS2, ALDH1B1, ALDH2, ALDOB, DLAT, DLD, ENO1, GPI, PDHA1, PDHB |
| Sulfur metabolism | 2.77e-05 | CYCS, ETHE1, MPST, SQRDL, TST |
| Synthesis and degradation of ketone bodies | 0.00051 | ACAT1, BDH1, HMGCL, HMGCS2 |
| Biosynthesis of unsaturated fatty acids | 0.00071 | ACOT1, ACOT2, ACOX1, HADHA, HSD17B12 |
| Fatty acid biosynthesis | 0.00086 | ACSL5, ACSL6, FASN, OXSM |
